# Intravital pH Imaging using the sensor ApHID Reveals Incomplete Lysosomal Acidification in Resident Meningeal Phagocytes

**DOI:** 10.64898/2026.09.12.750535

**Authors:** Santiago Solé-Domènech, Sung-Ji Ahn, Michael Iskols, Estibaliz Capetillo-Zarate, Lucy Funes, Jordi Pedragosa, Carles Justicia, Lynn-Marie Johnson, J David Warren, Anna Planas, Josef Anrather, Costantino Iadecola, Frederick R. Maxfield

## Abstract

Tracking lysosomes and quantifying their pH in the living, intact brain has been challenging owing to the lack of suitable in vivo tools. Here we report the use of dextran polymers labeled with the pH sensor ApHID to visualize and quantify lysosomal pH in resident phagocytes of the meninges. ApHID-dextrans delivered systemically are rapidly endocytosed by mainly dural and subdural macrophages and accumulate in lysosomal compartments. Intravital pH imaging revealed a striking heterogeneity in lysosomal acidification, with only ∼5% of phagocytes reaching pH values below 5.0, in marked contrast with measurements in macrophage cell cultures. Photothrombotic ischemic lesions induced progressive lysosomal acidification in local phagocytes, while acute chloroquine treatment led to rapid alkalinization. Our methodology enables direct, quantitative pH measurements without genetic manipulations, and unveils a previously unappreciated heterogeneity in lysosomal acidification among phagocytic cells in the meninges.

## INTRODUCTION

Lysosomes (Ly) are membrane-bound organelles containing more than 60 hydrolases and over 100 membrane proteins that constitute the degradative compartment of the endocytic system^1^. These organelles tightly regulate their intraluminal pH, which is essential for optimal hydrolytic activity^2–4^. Lysosomal enzymatic deficiencies lead to storage disorders such as Tay-Sachs disease and ceroid lipofuscinosis^5^. Lysosomal dysfunction has also been associated with aging^6^ and neurodegenerative diseases^7^. Intra-and extracellular, misfolded protein aggregation, including amyloid-β (Aβ), tau and α-synuclein, is a hallmark of Alzheimer’s disease (AD), Parkinson’s disease (PD) and tauopathies including frontotemporal dementia (FTD). Aggregates form intracellularly in part owing to declining lysosomal proteolytic capacity, and some subsequently accumulate as extracellular deposits^6^. Although microglia and leptomeningeal macrophages cluster around these lesions in the brain and leptomeningeal arterioles, aggregates persist and spread over time, causing neural damage as well as cognitive decline^8,9^. Some Ly proteases require pH ≤4.7 to efficiently proteolyze pathogenic fibrillar (f)Aβ, which is resistant to degradation owing to its β-sheet conformation^2,10^. However, the strongest genetic risk factors for late-onset AD –the ε4 allele of *APOE*^11^ and loss-of-function mutations in the myeloid receptor *TREM2*^12^– cause defective lysosomal acidification^13,14^, further involving lysosomes in pathogenesis.

Collectively, the involvement of lysosomes in aging and disease has been firmly established and some of the underlying mechanisms have been uncovered. However, these findings were obtained from cell cultures, while in vivo validation has remained challenging, owing to the lack of suitable intravital imaging tools. To address this, we used our recently developed pH sensor, ApHID^15^, conjugated to dextran polymers to measure endosomal pH in phagocytic cells *in the intact brain.* Dextrans have been used as *bona fide* markers of late endosomes and lysosomes (LE/Ly) for over 50 years in cell culture^16^. We injected these polymers systemically into mice, which led to internalization by meningeal phagocytes and accumulation into acidic vesicles, where they served as pH sensors. To monitor pH ratiometrically, in addition to ApHID, we also used the pH-independent probe Alexa Fluor 546, and imaged the meninges by two-photon microscopy through an implanted chronic cranial window.

We found that internalized dextrans accumulate intracellularly over time and adopt a perinuclear distribution consistent with LE/Ly compartments. Our ratiometric pH measurements revealed a substantial heterogeneity in LE/Ly acidification, with only 5% of all meningeal phagocytes acidifying their LE/Lys below pH 5.0. This is in sharp contrast with LE/Ly pH values established using macrophage cell culture models^4,17^. We also show that focal ischemic injury promoted LE/Ly acidification in local meningeal phagocytes, whereas systemic administration of chloroquine alkalinizes LE/Ly compartments within minutes.

To our knowledge, this is the first in vivo measurement of LE/Ly pH in phagocytes of the intact meninges. Our methodology can be used to label LE/Lys in any mouse model without requiring genetic manipulation, extending pH measurements to other contexts of aging and neurodegeneration and allowing exploration in an optimal, native biological environment.

## RESULTS

### ApHID fluorescence dynamic range matches the acidity of LE/Ly compartments and is resistant to photobleaching under two-photon excitation

Many commercially available fluorescent pH sensors have limited capacity to sense late endosomal and lysosomal (LE/Ly) pH (we will use LE/Ly rather than “lysosomes” because these are a collection of LAMP1-containing, acidic, hydrolytic organelles that exchange contents). For prolonged fluorescence excitation or intravital imaging of tissues, bright probes with robust fluorescence, resistance to chemical modification, and fluorescence dynamic range matching the acidity of acidic endosomes are needed. With that in mind, we designed a BODIPY-based pH-sensitive probe, called ApHID^15^ (Fig. 1A), a green-emitting sensor with a pKa of ∼5 (which falls at the center of the endosomal pH range) and fluorescence dynamic range optimally matched to the acidity of LE/Ly compartments. ApHID brightens with increasing acidity, resists photobleaching, and its fluorescence remains unchanged after paraformaldehyde (PFA) fixation^15^. We validated ApHID in solution and in cell culture extensively, demonstrating resistance to chemical and enzymatic degradation and photobleaching under one-photon excitation^15^. To investigate ApHID’s performance under two-photon (2-p) excitation, we derivatized 70 kDa amino dextran polymers with the probe and incubated J774A.1 macrophages with labeled dextrans overnight, followed by a 4h chase in fresh media (free of dextrans) to ensure LE/Ly localization. Endocytosed dextrans have been used as *bona fide* markers of LE/Ly compartments for over 50 years^16^. Cells were then mildly fixed in 0.5% PFA, and 50 mM TRIS maleate buffer adjusted to pH 5.5 was added at 37 °C in the presence of membrane permeant equilibrators. Images were acquired at several 2-p excitation wavelengths within the 690-1040 nm range (Fig. 1B). Fluorescence was quantified and plotted against excitation wavelength, rendering a qualitative 2-p excitation spectrum (Fig. 1C). We also tested ApHID-dextran’s resistance to 2-p photobleaching compared with fluorescein-dextran, a standard pH sensor used to measure LE/Ly pH^16^. Fixed J774A.1 macrophages labeled with the probes were incubated in pH 5.5 buffer at 37 °C and subjected to parallel 940 nm 2-p irradiation (50 images per stack, 0.26 s irradiation time per frame, 30 irradiation cycles). Laser output was adjusted to yield 50 mW power at the front element of the objective using an external power meter (Fig. 1D). Probe fluorescence was quantified and plotted against irradiation cycle, demonstrating ApHID’s resistance to 2-p photobleaching (Fig. 1E) in line with previous results under 1-p excitation^15^. Finally, to verify ApHID pH sensing under 2-p illumination, J774A.1 macrophages were loaded with 70 kDa dextrans tagged with ApHID and Alexa Fluor 546 (pH-independent), fixed and incubated in buffers with pH ranging 4.0-6.0 containing membrane-permeant equilibrators at 37 °C. Both probes were excited at 940 nm 2-p (Fig. 1F), and ApHID/Alexa 546 fluorescence ratios were plotted against buffer pH, rendering a curve that could be fit to a four-parameter sigmoid with IC_50_ of ∼5, matching ApHID-dextran pKa (Fig. 1G). Dextrans labeled with ApHID and Alexa Fluor 405 as a pH independent probe showed nearly identical pH-dependent dynamic range, validating this approach across multiple probes (Suppl. Fig. 1).

**Figure 1.**
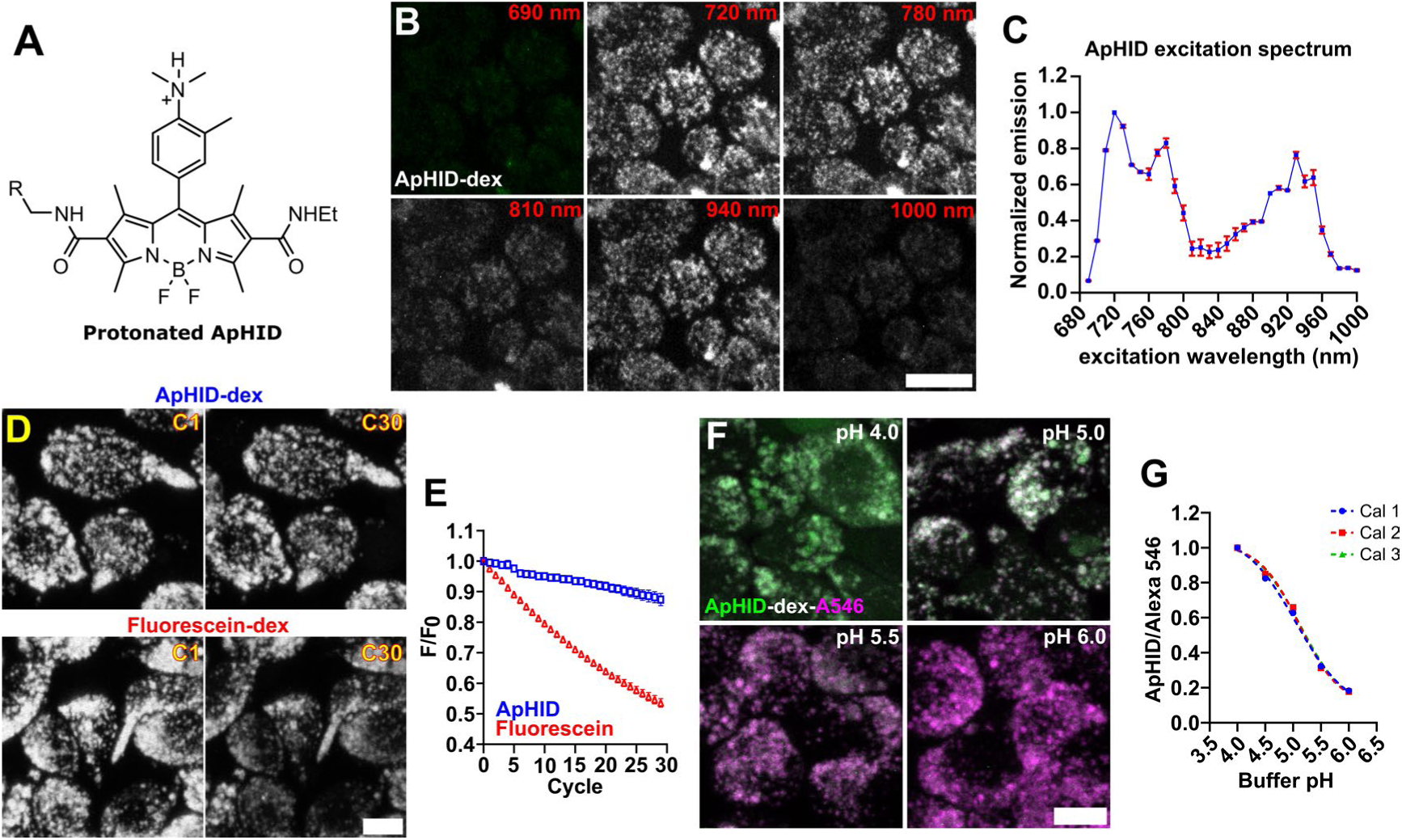
The pH sensor ApHID is two-photon excitable, resists photobleaching, and its fluorescence matches the endosomal acidic pH range. **(A)** Chemical structure of the fluorescent protonated ApHID molecule. **(B)** Fixed J774A.1 macrophages with late endosomes and lysosomes (LE/Ly) previously loaded with ApHID-dextrans (70 kDa), imaged in pH 5.5 buffer containing membrane-permeant equilibrators, under different two-photon excitation wavelengths (indicated in the top right corner of each panel, B). **(C)** ApHID fluorescence was quantified and plotted against two-photon excitation wavelength, yielding ApHID’s two-photon excitation spectrum. Fluorescence was normalized to 720 nm excitation. The experiment was repeated twice. 2 dishes were imaged in total and three fields were acquired per dish. Symbols and bars indicate mean normalized fluorescence ± SEM. **(D)** Fixed J774A.1 macrophages with LE/Lys loaded with ApHID-dextrans or fluorescein-dextrans. Cells were imaged in pH 5.5 buffer and irradiated with a two-photon laser tuned to 940 nm for 30 cycles. **(E)** Fluorescence in (D) was quantified and plotted against irradiation cycle for both probes. The experiment was repeated twice. 3-4 dishes were imaged per fluorophore and experiment, and three fields were acquired per dish. Scale bars: 20 µm. Symbols indicate mean fluorescence. Error bars (SEM) fit within the symbols. **(F)** Fixed J774A.1 macrophages labeled with dextrans conjugated with ApHID and Alexa 546 (pH-independent). Cells were incubated in buffers ranging from pH 4.0 to 6.0, containing membrane-permeant equilibrators, for 20–30 min and imaged at 37°C. **(G)** ApHID/Alexa 546 fluorescence ratios from images in (F) were plotted against buffer pH and fit to a four-parameter sigmoidal function, with an IC_50_ corresponding to ApHID’s pKa of ∼5. The experiment was repeated twice. One dish was imaged per pH condition, and four fields were acquired per dish. Symbols indicate mean fluorescence ratio. Bars (SEM) fit within the symbols. Scale bars: 20 µm (B, D); 10 µm (F). Abbreviations : dex : dextran ; A546 : Alexa 546.

Together, these findings indicate that ApHID can be excited by multiple 2-p excitation lines, is resistant to 2-p photobleaching, and maintains pH sensing capacity when excited using 2-p illumination. These properties turn ApHID into an optimal pH sensor for 2-p intravital imaging.

### Dextran sensors administered systemically into mice are rapidly endocytosed by meningeal phagocytes in vivo

To test dextran uptake by phagocytic cells in vivo, we focused on meningeal macrophages. We hypothesized that ApHID-dextran injected into the blood stream would be endocytosed by meningeal phagocytes in the dura, which is outside the blood-brain barrier (BBB) and accessible to circulating dextrans. The meninges were visualized intravitally through a cranial window in CX3CR1-eGFP mice, which express eGFP predominantly in meningeal macrophages^18^ (the majority of meningeal phagocytes in the dural and subdural compartments are macrophages^19,20^). We expected these phagocytes to robustly endocytose dextrans, as shown by various phagocytic cell lines in cell culture^15^. We injected CX3CR1-eGFP mice with dextrans intraperitoneally (i.p.) and imaged meningeal phagocytes intravitally shortly after injection (Fig. 2A). Prior to imaging, CX3CR1-eGFP^+^ cells appeared unremarkable with no detectable intracellular fluorescence, and the only detectable fluorescence corresponded to second-harmonic collagen signal (Fig. 2B-2C). However, following i.p. Alexa 405-dextran injection, these cells accumulated a strong intravesicular Alexa 405 signal (Fig. 2D-2E). CX3CR1-eGFP^+^ phagocytes close to the pia mater endocytosed dextrans too (Fig. 2F), but microglia sitting immediately underneath did not (Fig. 2G), consistent with the restriction to parenchymal penetrance imposed by the BBB on large molecules^21^. To quantify the fraction of cells that internalize dextran, we established a 5% labeled area threshold (Fig. 2H). Cells with their area occupied by ≥5% dextran were deemed positively labeled. 2h following initial i.p. injection, nearly two-thirds of all CX3CR1-eGFP^+^ phagocytes had internalized dextrans, whereas only 2% of labeled phagocytes were eGFP^-^ (readily identifiable by morphological criteria) (Fig. 2I).

**Figure 2.**
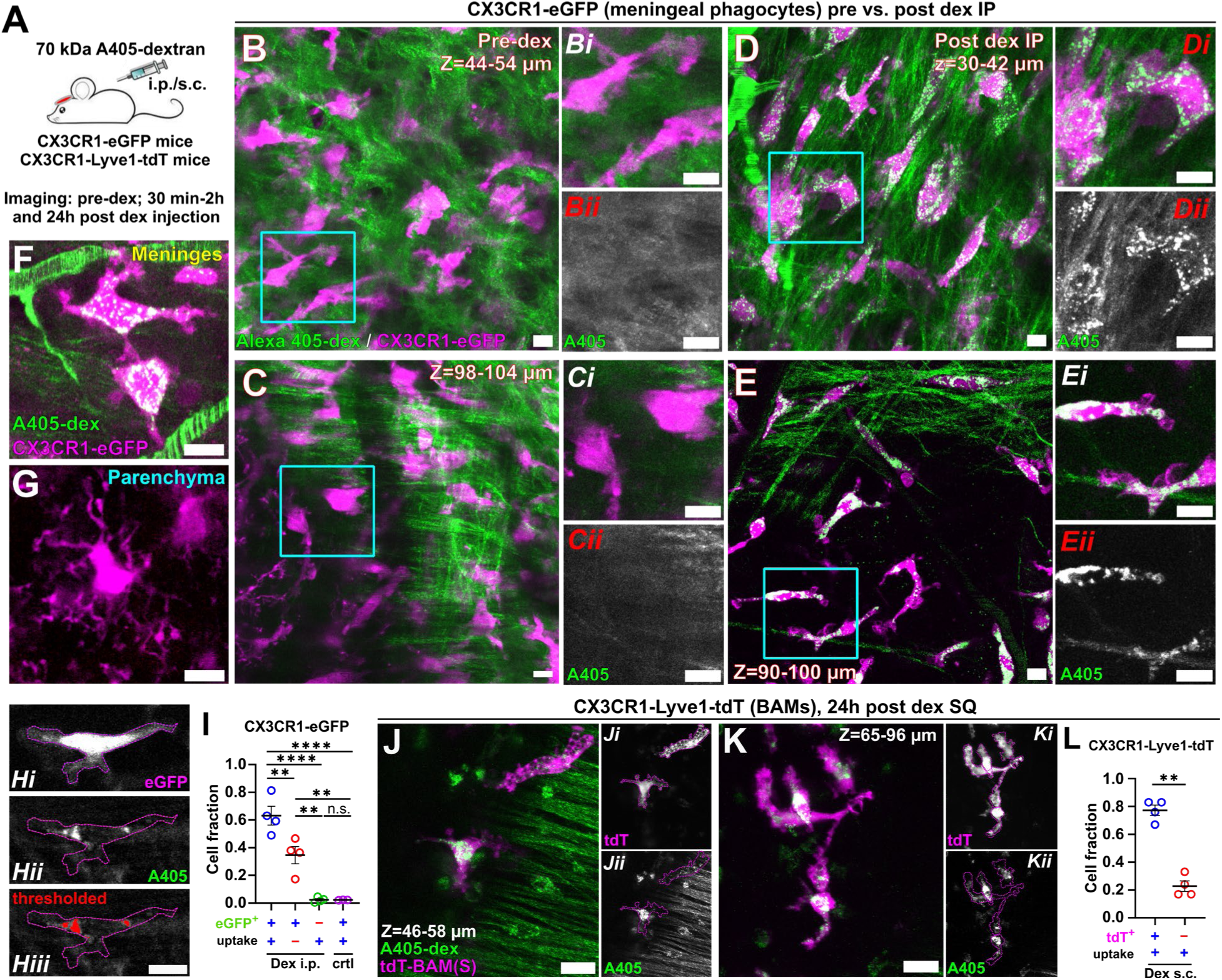
Dextrans injected intraperitoneally or subcutaneously are endocytosed by meningeal phagocytes. **(A)** Experimental strategy. Mice were implanted with chronic cranial windows and injected with dextrans systemically 2-3 weeks later. The meninges were visualized by intravital microscopy 30 min-2h (CX3CR1-eGFP mice) or 24h (CX3CR1-Lyve1-tdT mice) after dextran injection. **(B-C)** Representative images of eGFP^+^ phagocytic cells in the meninges, acquired at two depths prior to dextran delivery. Insets (cyan boxes) show individual eGFP⁺ phagocytes (Bi-Ci), and separate Alexa 405 channels (Bii-Cii) show filaments and background signal corresponding to second harmonic generation originating from collagen-rich dural layers. **(D-E)** Representative images of eGFP+ meningeal phagocytic cells at two depths, 30 min to 2h following dextran delivery (post-dex). Insets (cyan boxes) show individual eGFP⁺ phagocytes (Di-Ei), and separate Alexa 405 channels (Dii-Eii) show internalized Alexa 405-dextran signal, much brighter than second harmonic collagen signal and sufficiently salient in most cells to enable quantification in the meningeal space. **(F-G)** Representative images from a CX3CR1-eGFP mouse showing an eGFP⁺ (green) phagocyte in the subdural space (F) and a microglia in the adjacent cortical parenchyma (G), 30 min after dextran injection. Vasculature appears labeled with dextran (Alexa 405, red) in (F). **(H)** Representative example of a eGFP+ phagocyte (i) with internalized dextran (ii). Cells for which its thresholded Alexa 405 signal occupies ≥5% of the segmented cellular area (iii) were classified as positively labeled and quantified accordingly. **(I)** Alexa 405 dextran labeling was quantified in 1248 and 355 cells from 4 injected (‘Dex i.p.’) and 3 control (‘crt’) animals, respectively. Averaged fractions of GFP-positive and GFP-negative cells with or without dextran internalization (‘uptake’) between conditions were compared using a linear mixed effects model followed by Tukey’s multiple comparison test (p≥0.05 n.s.; p<0.01 **; p<0.0001 ****). **(J-K)** Representative images from a CX3CR1-Lyve1-tdTomato (tdT) mouse injected with 70 kDa Alexa 405-dextrans subcutaneously (24h post-dex), showing tdT^+^ macrophages acquired at two depths. Separate tdT (Ji, Ki) and Alexa 405-dextran (Jii, Kii) channels are shown. **(L)** Alexa 405 dextran uptake was quantified in 210 tdT^+^ dural and subdural macrophages from 4 injected animals. Averaged fractions of labeled and unlabeled cells were compared using paired Student’s t-test (p<0.01 **). In plots 2I and 2L, bars indicate mean cell fraction ±SEM per condition. Circles correspond to values for individual mice. Scale bars: large panels, 20 µm; insets, 10 µm. Abbreviations: A405: Alexa 405; A546: Alexa 546; dex: dextran; i.p.: intraperitoneally; s.c.: subcutaneously.

These cells could correspond to GFP-neutrophils, present in the dura and exhibiting phagocytic behavior^22^. Meningeal phagocytes in untreated mice did not show any remarkable intracellular fluorescence (Fig. 2I). This demonstrates that dextrans in the vasculature are internalized rapidly and selectively by CX3CR1-eGFP^+^ phagocytes in the meninges.

Since several leukocytes are CX3CR1^+^, we used the CX3CR1-Lyve1-tdTomato mouse line^23^ to more selectively image the internalization of dextran sensors by meningeal macrophages. CX3CR1 and Lyve1 promoters drive tdTomato expression selectively in a subset of meningeal and perivascular macrophages, and we used this system for cell-identity assignment. Mice implanted with cranial windows were injected with Alexa 405-labeled dextrans subcutaneously (s.c.) and allowed to recover for 24h followed by intravital imaging. 70 kDa dextrans are large biomolecules, and their absorption from the subcutaneous space is primarily mediated by the lymphatic network and occurs within minutes of injection. From the lymphatics, dextrans enter the bloodstream and reach meningeal vasculature^24^. 24h after s.c. delivery, dextrans had been completely absorbed (no subcutaneous liquid or bulge was present at the injection site), and no dextran fluorescence could be seen in meningeal vasculature. However, tdTomato^+^ macrophages appeared robustly labeled with Alexa 405-dextran (Fig. 2J-2K), with almost 80% showing intracellular dextran 24h post dextran injection (Fig. 2L).

Overall, these results indicate that meningeal phagocytes internalize dextran polymers injected into the blood stream and that this is a suitable system to test the ApHID-dextran pH probe.

### ApHID-dextran sensors can be used to measure endosomal pH intravitally in the intact meninges

In macrophage cell culture models, dextrans are internalized via fluid-phase pinocytosis (Fig. 3B) and trafficked into early endosomes (pH ∼6.0) within 3-10 minutes and LE/Ly compartments within 20-30 min (pH ∼5)^25^. To test this premise *in vivo* (Fig. 3A), we labeled 70 kDa dextrans with ApHID or fluorescein, and Alexa 546 (pH-independent), injected the polymers i.p. into mice and monitored dextran incorporation in the meninges intravitally (Fig. 3C). We observed robust fluorescein-Alexa 546-dextran internalization by dural and subdural meningeal phagocytes. The polymers accumulated in mildly acidic vesicles and in the vasculature, where fluorescein was bright (Fig. 3C). Fluorescein and Fluor 546 fluorescence was quantified per cell, in blood vessels (Fig. 3C) or in fixed macrophages (per field, Fig. 3D) and plotted. To do that, regions of interest were carefully drawn around the objects, avoiding overlap between perivascular cells and fluorescent, dextran-loaded vessels. Fluorescence in the pH-independent channel (Alexa 546) was thresholded and a mask was generated and applied to both the Alexa 546 channel and the pH-sensitive channel (fluorescein) thereafter.

**Figure 3.**
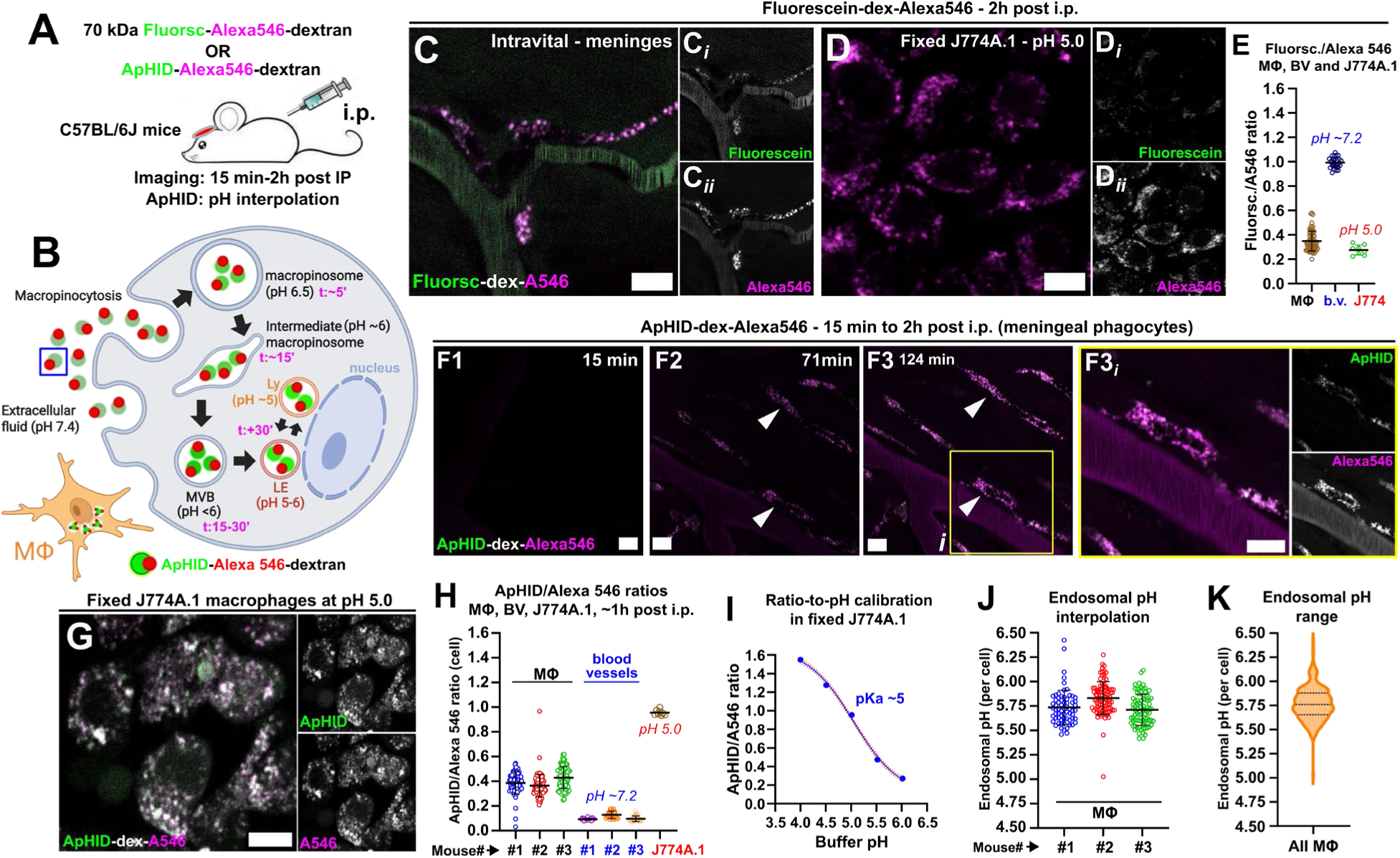
Ratiometric pH imaging of endosomal compartments in meningeal phagocytes using ApHID-dextran sensors. **(A)** Experimental strategy. Wild-type mice were implanted with chronic cranial windows and injected with dextrans intraperitoneally (i.p.) 2-3 weeks later. The meninges were continuously imaged by intravital microscopy 15 min after injection, for up to 2h. **(B)** Schematic representation of dextran internalization by fluid-phase pinocytosis as typically seen in phagocytic cells such as macrophages. Internalization times for each compartment are based on cell culture studies. **(C)** Intravital imaging of meningeal phagocytes (MΦ) in a C57BL/6J mouse injected with fluorescein-Alexa 546-dextrans (70 kDa). Dextrans incorporated into the vasculature, which also appears labeled. Side panels (i-ii) show individual fluorescence channels. **(D)** J774A.1 murine macrophages with late endosomes and lysosomes (LE/Lys) labeled with fluorescein-Alexa 546 dextrans (70 kDa), fixed in 0.5% PFA and incubated in pH 5.0 buffer containing membrane-permeant equilibrators at 37 °C. The cells were imaged alongside mice during the same intravital imaging session as a fixed-cell reference for calibration. Side panels (i-ii) show individual fluorescence channels. **(E)** Fluorescein/Alexa 546 fluorescence ratios calculated for individual meningeal phagocytes or individual blood vessels (n=1 mouse) and fixed J774A.1 macrophages in pH 5.0 buffer (D). **(F)** Representative intravital images of meningeal phagocytes in a C57BL/6J mouse injected i.p. with ApHID-Alexa 546-dextrans (70 kDa). Dextrans incorporated into the vasculature, which also appears labeled. Images were acquired from the same region at various times post-injection (F1-F3). The inset shows individual phagocytes with abundantly labeled acidic vesicles (F3i). Side panels show individual fluorescence channels. **(G)** J774A.1 murine macrophages labeled with ApHID-Alexa 546 dextrans (70 kDa), fixed and incubated in pH 5.0 buffer at 37 °C, imaged immediately following intravital imaging as described in (D). **(H)** ApHID/Alexa 546 fluorescence ratios calculated for individual meningeal phagocytes and blood vessels at 70-125 min post-i.p. injection time (F2-F3, n=3 mice, shown individually) or for fixed J774A.1 macrophages (ratios per field) in pH 5.0 buffer at 37 °C (G). **(I)** ApHID/Alexa 546 ratios measured in fixed macrophages in pH 5.0 buffer (G) were used to generate a ratio-to-pH standard calibration curve, with IC_50_ corresponding to ApHID’s pKa (∼5), as previously described^15^. One dish was incubated in pH 5.0 buffer for 36 min at 37 °C and imaged thereafter (8 fields acquired) at the end of each technical replicate. Symbols indicate mean ratio per pH point. Error bars (SEM) fit within the symbols. The dotted curve corresponds to the sigmoidal fit. **(J-K)** ApHID/Alexa 546 fluorescence ratios for each segmented meningeal phagocyte (MΦ) were translated to pH by interpolation to a ratio-to-pH calibration (as in I) and plotted per mouse (J) or pooled together (K). In plots (E), (H) and (J), symbols correspond to values for individual meningeal phagocytes, blood vessels or fields containing fixed J774A.1 macrophages. Bars indicate mean ratio or pH ±SD. In (K), dotted lines indicate median and quartiles. Scale bars: 10 µm. Abbreviations: MΦ: meningeal phagocytes; dex: dextran; b.v.: blood vessels; i.p.: intraperitoneally.

Fluorescence in the masked channels was quantified for each drawn region, and fluorescence ratios for each object were calculated. Ratios measured for blood vessels were highest, corresponding to increasing fluorescein signal with increasing pH, whereas ratios measured in fixed J774A.1 macrophages at pH 5.0 (Fig. 3D) were the lowest, consistent with fluorescein decreasing brightness with acidity. Fluorescein/Alexa 546 ratios measured in real time in meningeal phagocytes suggest a pH >5.0 (Fig. 3E).

ApHID-dextran pKa is ∼5, rendering the probe more sensitive in the endosomal pH range relative to fluorescein (pKa ∼6.4)^15^. To test ApHID capacity to report pH in real time, we labeled dextrans with it and Alexa 546 and injected them i.p. into mice as described above. Meningeal vasculature progressively incorporated the polymers (Fig. 3F), and meningeal phagocytes gained sufficient intracellular signal for quantitative measurements 70 min after injection (Fig. 3F2, arrowheads). ApHID signal was weak in the vasculature, corresponding with its decreasing fluorescence with alkalinity. As a reference, fixed J774A.1 macrophages loaded with the same dextrans and imaged in pH 5.0 buffer showed brighter ApHID signal (Fig. 3G).

ApHID/Alexa 546 fluorescence ratios calculated for meningeal phagocytes, blood vessels and fixed macrophages reflected ApHID’s increasing brightness with acidity (Fig. 3H). In order to interpolate endosomal fluorescence ratios to pH, a buffer-to-pH standard calibration was built using ApHID/Alexa 546 fluorescence ratio at pH 5.0 from fixed cells (Figs. 3D), and the remaining ratio values were generated using complete titration data prepared previously using fixed J774A.1 macrophages imaged with the same two-photon microscope (Fig. 3I). This approach is possible because ApHID dynamic range is stable at a given temperature (in our case, 37 °C) and PFA fixation does not alter its fluorescence properties^15^. pH was calculated for individual meningeal phagocytes (n=3 mice) and plotted per mouse (Fig. 3J) or pooled for all cells (Fig. 3K), resulting in a mean endosomal pH of 5.75. This reflects a mixed endosomal population, composed of early and late endosomes, which is consistent with the imaging time frame (∼1h post IP, Fig. 3B). Importantly, fluorescein/Alexa 546 ratios remained stable within dural and subdural compartment depths –fluorescein shares a near-identical emission profile with ApHID but fluoresces at neutral blood pH, serving as a light-scattering reference (Suppl. Fig. 2). This demonstrates minimal or no differential optical aberration by depth-associated light scattering on probe fluorescence.

Overall, our ratiometric pH imaging of endosomes in meningeal phagocytes demonstrate that ApHID can be used to quantify endosomal pH in vivo in the intact meninges.

### Resident meningeal phagocytes do not fully acidify late endosomal and lysosomal compartments

In phagocytic cells in tissue culture, dextrans are endocytosed by fluid phase pinocytosis and trafficked to LE/Lys within 20-30 minutes of initial internalization^25^, and a 3-4h chase in fresh medium ensures dextran localization in these organelles^15^. We asked whether a similar strategy could be applied to label LE/Ly compartments and measure their acidity intravitally. LE/Ly pH measured in macrophage cell lines using ratiometric pH imaging ranges between pH 4.5 and 5.0^4^. We define LE/Ly compartments as fully acidified when they reach pH≤4.7, the threshold at which the combined activity of lysosomal proteases becomes maximal^10,26^. This can enhance degradation of pathogenic substrates such as amyloid-beta or tau fibrils, which are resistant to degradation due to their beta-sheet conformation^2^. Building on this premise, we next asked whether resident meningeal macrophages achieve the same degree of LE/Ly acidification in vivo.

We injected CX3CR1-Lyve1-tdTomato mice with 70 kDa polymers labeled with Alexa 405 delivered s.c. followed by a 24h recovery period (Fig. 4A). Intravital imaging revealed a mainly perinuclear dextran distribution in tdT+ macrophages, consistent with the classic LE/Ly localization seen in cell culture models (Fig. 4B). Dextrans labeled with ApHID and Alexa 546 injected s.c. were also taken up by meningeal phagocytes, which showed a ∼100-fold increase in intracellular green and red fluorescence 24h after injection relative to pre-injection condition (Fig. 4C), demonstrating robust uptake and optimal signal for quantitative purposes.

**Figure 4.**
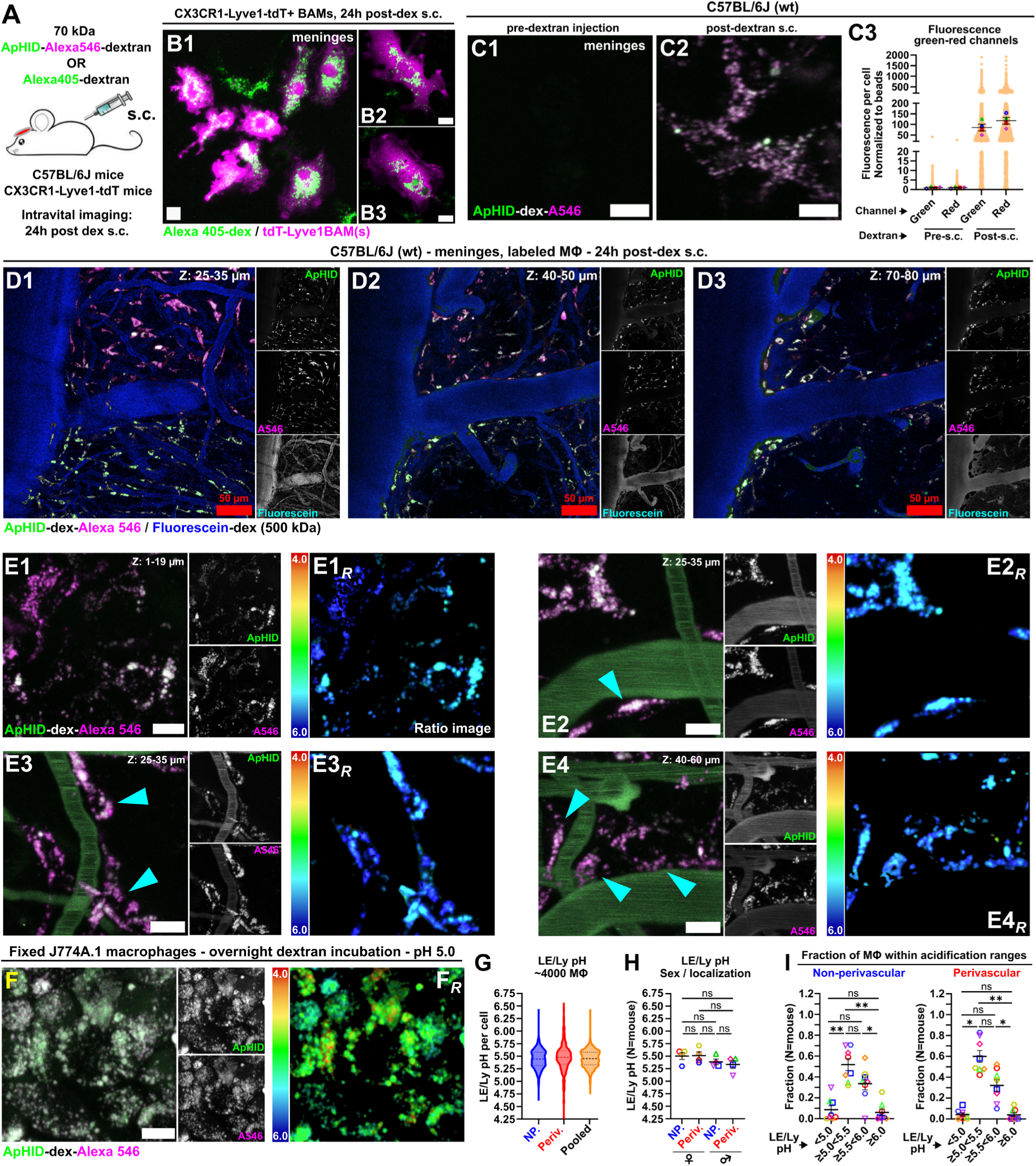
Meningeal phagocytes do not fully acidify their late endosomes and lysosomes. **(A)** Experimental strategy. Wild-type and mutant CX3CR1-Lyve1-tdT mice were implanted with chronic cranial windows and injected with dextrans subcutaneously (s.c.) 2-3 weeks later. The meninges were imaged by intravital microscopy 24h later. **(B)** Intravital imaging of meningeal tdT+ macrophages in a CX3CR1-Lyve1-tdT mouse injected with Alexa 405-dextrans (70 kDa) and allowed to recover for 24h prior to imaging. Dextrans show a perinuclear distribution consistent with late endosomal and lysosomal localization. **(C)** Intravital imaging of meningeal phagocytes in a C57BL/6J mouse prior to (C1) or 24h after (C2) s.c. injection with ApHID-Alexa 546-dextrans (70 kDa). Fluorescence values in meningeal phagocytes for ApHID (green channel) and Alexa 546 (red channel) measured before (Pre-s,c.) or after (Post-s.c.) dextran injection are plotted normalized to the average Pre-s.c. value (C3). To control for variations in laser power between imaging sessions, fluorescence values were additionally normalized to Nile red bead fluorescence recorded at the end of each session. Large symbols indicate integrated intensity values per mouse (n=4) and small tan circles correspond to individual segmented phagocytes. Bars indicate average fluorescence intensity ± SEM. **(D)** Intravital pH imaging of acidic late endosomal and lysosomal (LE/Ly) compartments in meningeal phagocytes acquired through a cranial window in a C57BL/6J mouse. The animal was injected s.c. with ApHID-Alexa 546 dextrans (70 kDa) and allowed to recover for 24h. Vasculature was labeled by intraperitoneal (i.p.) injection of fluorescein-Alexa 546 dextrans (500 kDa) 10 min prior to imaging. The same region was imaged at various depths (D1-D3). **(E)** Intravital pH imaging of meningeal phagocytes as described in (D), showing cells at the surface of the dura (E1) and others immediately adjacent to blood vessels in deeper meningeal layers (E2-E4, perivascular localization, arrowheads). Vasculature was labeled with fluorescein-Alexa 546 dextrans (500 kDa) 10 min prior to imaging. Color-coded ratiometric pH images were generated for each region (E1R-E4R). Color-coded bars indicate a pH range of 4.0 (red) to 6.0 (blue). **(F)** J774A.1 murine macrophages labeled with ApHID-Alexa 546 dextrans (70 kDa), fixed and incubated in pH 5.0 buffer at 37 °C, imaged immediately following intravital imaging. A color-coded ratiometric pH image was generated as described in (E). **(G)** ApHID/Alexa 546 fluorescence ratios corresponding to late endosomal and lysosomal (LE/Ly) compartments for each segmented meningeal phagocyte (D-E) were interpolated to pH using a ratio-to-pH calibration (as shown in Fig. 3I). Violin plots show integrated LE/Ly pH including all segmented phagocytes (4093 cells) from 8 mice (4 males and 4 females). Cells in close proximity to blood vessels (“Periv.”) or further from vasculature (“NP”, generally at the dural surface) were plotted separately. Superimposed dotted lines indicate median and quartiles. **(H)** Mean LE/Ly pH (n=8, 4 males and 4 females) plotted according to vascular localization (“perivascular” vs “non-perivascular”) and sex. Mean LE/Ly pH between male and female mice and cells sitting in close proximity or away from vasculature were compared using a linear mixed effects model followed by Tukey’s multiple comparison test (p≥0.05 n.s.; all comparisons). Bars indicate mean LE/Ly pH ±SEM whereas individual symbols correspond to values for individual mice. **(I)** Meningeal phagocytes plotted against their LE/Ly pH range (pH<5.0; pH≥5.0–<5.5; pH≥5.5–<6.0; pH≥6.0). The mean fraction of cells within each range was calculated per mouse (n=8, 4 males and 4 females) and plotted by proximity to blood vessels (perivascular vs. non-perivascular) as shown in (E). Differences in cell fraction mean ranks were assessed using the non-parametric paired Friedman test with multiple comparisons by Dunn’s test (p≥0.05 n.s.; p<0.05 *; p<0.01 **). Bars indicate mean fraction of cells ±SEM whereas symbols correspond to values for individual mice. Side panels in D, E and F show individual fluorescence channels. Scale bars: Panels (D1-D3): 50 µm. All other panels: 10 µm. Abbreviations: A405: Alexa 405; A546: Alexa 546; dex: dextran; s.c.: subcutaneously; i.p.: intraperitoneally; MΦ: meningeal phagocytes.

Intravital ratiometric pH imaging of the meninges revealed striking heterogeneity in phagocyte acidification. Adjacent phagocytes within the same meningeal region often showed sharply different LE/Ly pH values at steady state, despite no evidence of vascular or tissue alterations that could impair cellular ability to sustain LE/Ly acidification. This pattern was consistent across several fields and depths imaged (Fig. 4D). Vasculature was labeled with 500 kDa fluorescein-dextrans, which are not internalized by phagocytes within our imaging times (Suppl. Fig. 3). Meningeal phagocytes showed robust dextran uptake (Fig. 4E1–4E4, arrowheads), and color-coded pH imaging confirmed pH gradients between cells and within individual cells (Fig. 4E1R–4E4R).

As a reference, fixed J774A.1 macrophages loaded with the same dextrans and imaged in pH 5.0 buffer showed brighter ApHID signal (Fig. 4F–4FR). ApHID/Alexa 546 fluorescence ratios quantified for more than 4,000 meningeal phagocytes (imaged across 4 male and 4 female wild-type mice) were converted to pH by interpolation using a ratio-to-pH standard curve (as in Figs. 1G and 3I), revealing a wide acidification range (pH 4.75-6.5) which seemed independent of phagocyte proximity to vasculature (Fig. 4G-4H). There was no difference in LE/Ly acidification between male and female mice (Fig. 4H). About half of all meningeal phagocytes showed LE/Ly pH values ranging between 5.0-5.5, and one-third of the cells fell between pH 5.5 and 6.0 (Fig. 4I). A small fraction of phagocytes showed LE/Ly pH above 6.0. Strikingly, only ∼5% of all phagocytes acidified their LE/Lys below pH 5.0. This is in marked contrast with studies in cell culture models, in which most macrophage lines exhibit LE/Ly pH values between 4.5-5.0^4,15^.

Overall, our data indicate that dextran polymers delivered s.c. in mice accumulate in intracellular acidic vesicles with a distribution consistent with LE/Ly compartments. pH measurements using ApHID-dextrans revealed a marked heterogeneity in meningeal phagocyte acidification, with only a small fraction of cells acidifying LE/Lys below pH 5.0.

### Lysosomal acidification in meningeal phagocytes is not dependent on proximity to arterioles or venules

We asked whether proximity of meningeal phagocytes to arterioles or venules, under non-pathological conditions, determines LE/Ly acidification. To test this, we injected wild-type mice with ApHID-Alexa 546 dextrans delivered s.c. and imaged their meninges intravitally 24h later. Immediately prior to imaging, we labeled meningeal vasculature by retro-orbital injection of 500 kDa Alexa 405 dextrans, which allowed us to identify and discriminate arterioles from venules based on the direction of flow (Fig. 5A). Penetrating arterioles were distinguished by the presence of vasomotion^27^ and inward blood flow into cortical parenchyma^28^, whereas venules had larger lumens and drained toward the cortical surface (Fig. 5B). We acquired multiple fields containing dextran-loaded meningeal phagocytes in close proximity to arterioles or venules (Fig. 5C–5D). As a reference, fixed J774A.1 macrophages loaded with the same dextrans were imaged in pH 5.0 buffer, as described above (Fig. 5E). ApHID/Alexa 546 fluorescence ratios were interpolated to pH as described earlier and plotted against proximity to arterioles or venules, revealing no difference in LE/Ly acidification (n=3 mice, 211 cells quantified, Fig. 5F).

**Figure 5.**
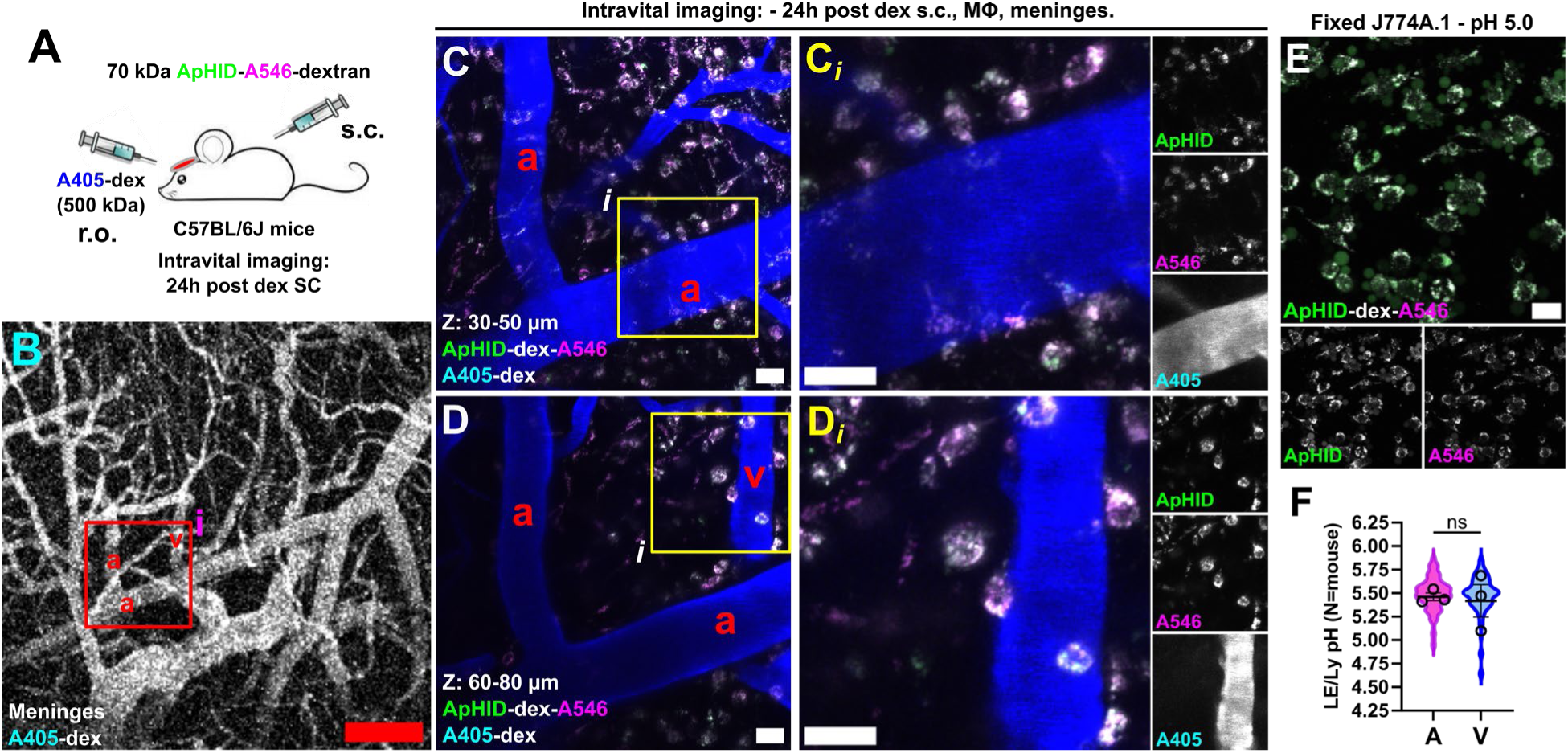
Meningeal phagocyte proximity to venules or arterioles does not influence lysosomal pH. **(A)** Experimental strategy. Wild-type mice were implanted with chronic cranial windows and injected with 70 kDa dextrans systemically 2-3 weeks later. The meninges were imaged by intravital microscopy 24h later. The vasculature was loaded by r.o. injection of 500 kDa Alexa 405-dextrans immediately prior to imaging. **(B)** Intravital imaging of meningeal vasculature labeled with Alexa 405-dextran (500 kDa) delivered retro-orbitally (r.o.) prior to imaging. The inset (i) highlights an arteriole (a) and a venule (v). **(C-D)** Expanded inset (Bi) imaged at two depths (C and D). Yellow squares (i) delineate an arteriole (C) and a venule (D), further expanded in the adjacent panels (Ci and Di). Side panels show individual fluorescence channels. **(E)** J774A.1 murine macrophages labeled with ApHID-Alexa 546 dextrans (70 kDa), fixed and incubated in pH 5.0 buffer at 37 °C, imaged immediately following intravital imaging. Side panels show individual fluorescence channels. **(F)** Mean LE/Ly pH (n=3 mice) in meningeal phagocytes, plotted by proximity to arterioles (A) or venules (V). Violin plots integrate LE/Ly pH for all segmented phagocytes (211 cells). Bars indicate mean LE/Ly pH ± SEM whereas symbols correspond to values for individual mice. Mean LE/Ly pH between conditions was compared using the paired Student’s t test (p≥0.05 n.s.). Scale bars: B: 200 µm; C, D: 20 µm. A405: Alexa 405; A546: Alexa 546; dex: dextran; MΦ: meningeal phagocytes; A546; s.c.: subcutaneously.

These data suggest that, under non-pathological conditions, meningeal phagocyte proximity to arterioles or venules does not influence LE/Ly acidification.

### Focal ischemic injury induces lysosomal acidification in local phagocytes

As mentioned previously, a significant fraction of phagocytes that internalize dextrans in the meninges are macrophages. These cells typically respond to inflammation by upregulating phagocytic activity and lysosomal acidification, which facilitates clearance and degradation of pathogens and cellular debris^29,30^. Evidence of this has been obtained using macrophage cell culture models and biological tissues, but LE/Ly pH has never been quantified in active phagocytes of the intact meninges under pro-inflammatory activation. To address this, we induced local ischemic lesions –a known inflammatory insult^31^– by photothrombosis of single penetrating arterioles loaded with Rose Bengal^31^ and monitored LE/Ly pH over time in local phagocytes localized to the lesioned areas.

Fig. 6A summarizes the experimental paradigm. Prior to each imaging session, 5–6-week-old mice were injected s.c. with 70 kDa amino-dextrans labeled with ApHID and Alexa 546 and housed overnight. To induce photothrombosis, mice were injected with Rose Bengal^32^ and penetrating arterioles were selected for targeted laser illumination. Mice were imaged intravitally at multiple time points following ischemic lesion. Images of selected fields were acquired before and after photothrombosis. Mice were also injected with 500 kDa Alexa 405-dextrans retro-orbitally immediately prior to each imaging session to label vasculature and relocate the same lesioned regions consistently across sessions. A representative brain region 11 days after lesion induction is shown in Fig. 6B and Fig. 6C. Blood flow was interrupted immediately after 720 nm illumination (Fig. 6E), but all targeted arterioles had spontaneously recanalized by the first follow-up session, evidenced by intraluminal filling with 500 kDa Alexa 405-dextran and visible red blood cell flow. This transient occlusion is consistent with the present measurements, since ApHID-dextran sensors are delivered systemically at each imaging session and require intact blood flow to reach phagocytes within the lesioned area. An occluded vessel would prevent sensor delivery entirely.

**Figure 6.**
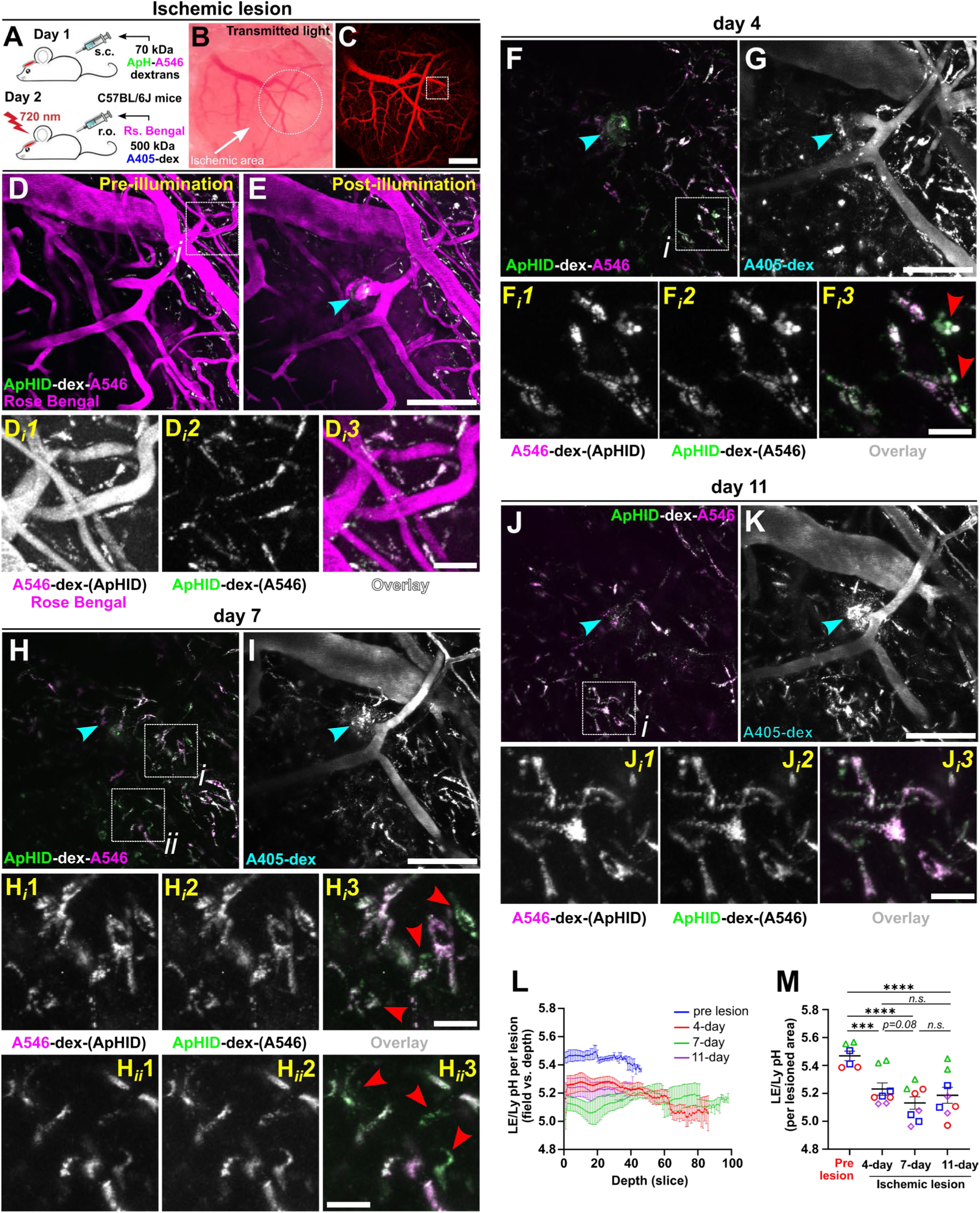
Meningeal ischemic insult promotes lysosomal acidification in nearby phagocytic cells. **(A)** Experimental strategy. Wild-type mice were implanted with chronic cranial windows and injected with 70 kDa dextrans systemically 2-3 weeks later. The meninges were imaged by intravital microscopy 24h later. Rose Bengal was injected retro-orbitally (r.o.) immediately prior to photothrombosis, induced by two-photon 720 nm laser illumination of single meningeal penetrating arterioles. Mice were re-imaged at 4, 7 and 11 days following focal ischemic lesion, and injected with 70 kDa dextrans systemically the day before imaging. The vasculature was loaded by r.o. injection of 500 kDa Alexa 405-dextrans immediately prior to each imaging session. **(B)** Transmitted-light image showing the ischemic area 11 days after initial photothrombosis, with the lesioned area outlined by a dotted white circle. **(C)** Two-photon wide-field image of the cortical vasculature. The white square indicates the location of the higher magnification image shown in D. **(D)** Cortical vasculature prior to photothrombotic laser illumination in a mouse in which Rose Bengal was injected retro-orbitally. 70 kDa amino dextran polymers (ApHID-dex-Alexa 546 (A546)) were injected a day prior. The inset (white square, i) highlights phagocytic cells with intracellular dextran labeling. The magnified views show Alexa 546 and Rose Bengal (Di1), ApHID (Di2), and their overlay (Di3). **(E)** Same field as in (D) immediately after photothrombosis induced by photoactivation of Rose Bengal with 720 nm laser light. Arrowheads highlight the lesion site. **(F)** Follow-up image of the same field 4 days post-photothrombosis. The inset (white square, i) highlights magnified views of A546 (Fi1), ApHID (Fi2), and their overlay (Fi3). Red arrows indicate acidified cells. **(G)** Same area as in (F), showing cortical vasculature labeled with 500 kDa Alexa 405-dextrans injected retro-orbitally (r.o.) immediately prior to imaging. Phagocytic cells near the lesion site internalized A405-dextrans, suggesting increased blood–brain barrier permeability after the insult. **(H–I)** Same area as in (F), 7 days post-photothrombosis. Insets highlight regions (i, ii), with magnified views showing A546 (Hi1-Hii1), ApHID (Hi2-Hii2) and their overlay (Hi3-Hii3). Cells remain acidified, and a stronger Alexa 405 signal is observed near the targeted vessel (I). **(J–K)** Same area as in (F), 11 days post-photothrombosis. The inset (i) highlights magnified views showing A546 (Ji1), ApHID (Ji2) and their overlay (Ji3). Cells remain acidified, with persistent Alexa 405 signal near the targeted vessel (K). **(L)** Mean meningeal phagocyte LE/Ly pH in lesioned areas (1-2 brain regions acquired per animal, n=4 mice) imaged over time, for each optical plane and plotted against imaging depth. Stacks were acquired at 1-µm axial steps. Bars indicate mean LE/Ly pH ±SEM. **(M)** To assess acidification across the entire lesion independent of depth or cell type, sum projections were generated from each stack, and ApHID/Alexa 546 ratios were converted to pH. Bars indicate average LE/Ly pH over time ±SEM (1-2 brain regions acquired per animal, n=4 mice). Symbols correspond to values for individual lesioned areas. Differences in LE/Ly pH means between time points were compared using a linear mixed model followed by Tukey’s multiple comparison test (n: lesioned area; p≥0.05 n.s.; p<0.001 ***; p<0.0001 ****). Scale bars: large panels: 500 µm (panel C), 100 µm (all other panels); insets: 20 µm. Abbreviations: dex: dextran; A546: Alexa 546; A405: Alexa 405; s.c.: subcutaneously; r.o.: retro-orbitally; Rs. Bengal: Rose Bengal.

A meningeal arteriole prior to and immediately after photothrombotic lesion induction is shown in Figures 6D and 6E. Following 720 nm laser illumination, blood flow appeared interrupted in the targeted arteriole (Fig. 6E), demonstrating successful clotting. Local phagocytic cells imaged prior to lesion induction showed robust ApHID–Alexa 546 dextran internalization (Fig. 6D, insets 6Di1–6Di3). The lesioned area was subsequently reimaged 4 days (Fig. 6F–6G), 7 days (Fig. 6H–6I), and 11 days (Fig. 6J–6K) after photothrombosis.

Phagocyte recruitment was observed around lesioned arterioles at 7 and 11 days post-lesion (Fig. 6H and 6K). These recruited phagocytes had internalized 500 kDa Alexa 405-dextrans (which do not interfere with pH measurements, Fig. 6G, 6I, 6K), suggesting disruption of vascular integrity and barrier function, as 500 kDa dextrans are generally too large to penetrate the meninges under basal conditions (Suppl. Fig. 3). Some meningeal phagocytes transitioned from an elongated morphology pre-lesion (Fig. 6D–E) to amoeboid morphologies post-lesion (Fig. 6Fi1–6Fi3, 6Hi1–6Hii3, and 6Ji1–6Ji3). ApHID and Alexa 546 fluorescence was quantified per optical plane, or as sum projections per stack, and ApHID/Alexa 546 ratios were calculated and interpolated to pH. LE/Ly pH plotted against imaging depth (Fig. 6L) or post-lesion time (Fig. 6M) for each lesioned region revealed a progressive acidification (from baseline pH at ∼5.3) following ischemic lesion, reaching a plateau at 7-11 days post-lesion (Fig. 6M). 7 days following photothrombosis, some regions showed a pH drop of up to ∼0.5 pH units relative to baseline. At 7 and 11 days post-lesion, dextran-loaded cells with rounded morphologies were observed at 40–100 µm depths within the lesioned areas, suggesting macrophage recruitment from the periphery (Suppl. Fig. 4).

Overall, these findings suggest that focal ischemic lesions–a well-documented inflammatory insult^33^– induce progressive LE/Ly acidification and morphological changes in local meningeal phagocytes. Infiltrating phagocytes with more acidic LE/Lys were observed within lesioned areas at 7 and 11 days post-lesion. This demonstrates that ApHID-dextrans can be used to monitor changes in LE/Ly acidification in real time under pro-inflammatory conditions.

### Chloroquine alkalinizes lysosomal pH within minutes in meningeal phagocytes

To test whether we can detect LE/Ly alkalinization in vivo using ApHID, we measured pH in dural phagocytes of mice treated acutely with chloroquine diphosphate (CQ). CQ is a weak base that crosses biomembranes and accumulates in endosomes, causing alkalinization^34^. We injected mice i.p. with a single dose of chloroquine diphosphate (500 mg/Kg) sufficient to drive rapid endosomal alkalinization in mice^35^ (Fig. 7A). Dural phagocytes with acidic LE/Ly compartments (Fig. 7B1 and insets) underwent rapid alkalinization 2 min (Fig. 7B2 and insets) and 8 min (Fig. 7B3 and insets) after CQ injection, demonstrated by a sharp drop in ApHID fluorescence (Fig. 7B1i–7B3i, color-coded ratiometric pH images), whereas fixed J774A.1 macrophages incubated in pH 5.0 buffer showed robust, stable ApHID fluorescence (Fig. 7C). pH interpolation indicated a LE/Ly pH increase in dural phagocytes of ∼0.4 pH units within 8 min following CQ injection (Fig. 7D), a value well within ApHID’s reliable measuring range.

**Figure 7.**
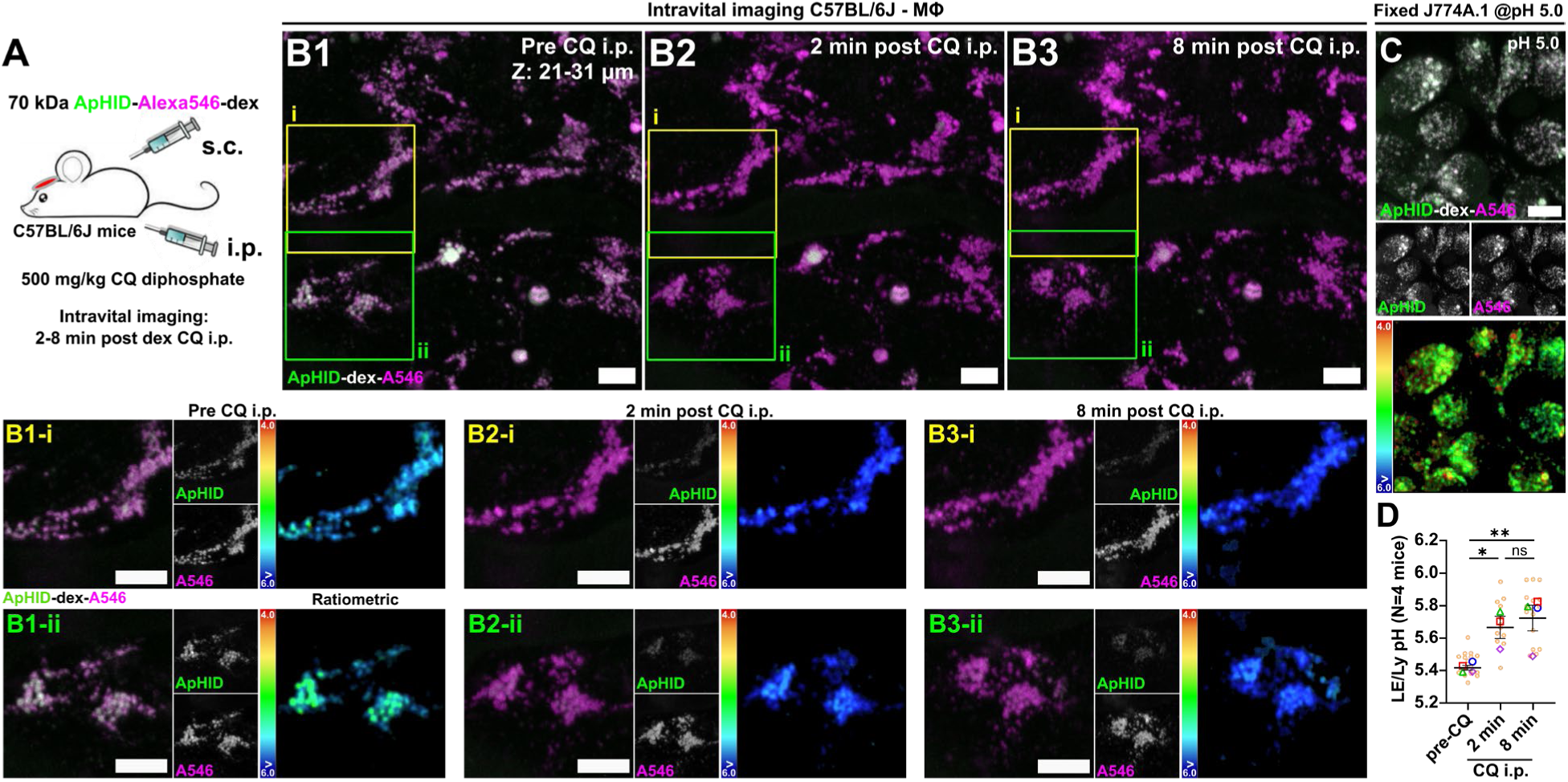
**Chloroquine rapidly alkalinizes late endosomal and lysosomal compartments in meningeal phagocytes.** (A) Experimental strategy. Wild-type mice were implanted with chronic cranial windows and injected with 70 kDa dextrans systemically 2-3 weeks later. The meninges were imaged by intravital microscopy 24h later. Immediately prior to imaging, mice were injected i.p. with an acute dose of chloroquine diphosphate (500 mg/Kg). (B) Intravital pH imaging of meningeal phagocytes before (B1) or 2 min (B2) or 8 min (B3) after chloroquine diphosphate i.p. delivery. Squares (i, ii) delineate two phagocytes before and after CQ delivery, with insets expanded in B1-i/ii, B2-i/ii, and B3-i/ii. Ratiometric images were generated for each expanded inset. Color-coded bars indicate pH range: 4.0 (red) to >6.0 (blue). Side panels in B1-i through B3-ii show individual fluorescence channels. (C) J774A.1 murine macrophages labeled with ApHID-Alexa 546 dextrans (70 kDa), fixed and incubated in pH 5.0 buffer at 37 °C, imaged immediately following intravital imaging. A color-coded ratiometric pH image was generated as described in (B). (D) Mean LE/Ly pH for meningeal phagocytes calculated across entire acquired stacks and averaged per mouse (n=4, 3 females and 1 male), plotted against post-CQ injection time. Bars indicate mean LE/Ly pH ± SEM. Large and small symbols correspond to values for individual mice or imaged fields within mice, respectively. Mean LE/Ly between timepoints was compared using a linear mixed model followed by Tukey’s multiple comparison test (p≥0.05 n.s.; p<0.05 *; p<0.01 **). Scale bars: 10 µm. A546: Alexa 546; CQ: chloroquine diphosphate; dex: dextran; MΦ: meningeal phagocytes; s.c.: subcutaneously; r.o.: retro-orbitally.

Overall, these data demonstrate that ApHID can accurately report alkalinization of LE/Ly pH in vivo and in real time in an intact meninges.

## DISCUSSION

Some important studies reported the use of mouse models to track macroautophagy or mitophagy with fluorescent markers^36–38^. Maeda and collaborators used pH imaging to measure pH of osteoclast lacunae in real time in mice^39^, and Kim and collaborators measured lysosomal acidification associated with autophagy in tumor environments using near-infrared optical carbon nanotubes^40^. In our study, we used amino-dextran polymers, a bona fide marker of LE/Ly compartments used for over 50 years^16^, to label and image acidic compartments in an intact meninges. To our knowledge, this is the first time LE/Ly compartments are labeled and their acidity measured quantitatively *in vivo* in meningeal phagocytes of wild-type mice.

In our hands, dextrans injected systemically (via i.p. or s.c. routes) were rapidly internalized by dural and subdural phagocytic cells. In CX3CR1-eGFP mice, eGFP is expressed in macrophages, natural killer cells, T cells and a fraction of dendritic cells in the meninges, and by microglia in the parenchyma. We imaged through dural and subdural spaces, and these compartments contain a heterogeneous immune population, including neutrophils, various subsets of monocytes and dendritic cells, NK cells, B cells and BAMs^19^. While we refer to dural and subdural dextran-labeled cells as meningeal phagocytes throughout, the predominant phagocytic cell type in the imaged compartments corresponds to macrophages^19,20^. The robust dextran uptake by CX3CR1-Lyve1-tdT+ dural and subdural macrophages is consistent with this premise^30^. Only ∼2% of meningeal phagocytes containing intracellular dextran were eGFP^-^, which could correspond to phagocytic neutrophils^22^.

Dextran polymers accumulated in acidic vesicles, leading to very bright, quantifiable fluorescence. Once endocytosed, and if labeled with pH-sensitive probes, dextrans can be used as pH sensors. We used this property to measure endosomal pH in meningeal phagocytes shortly after dextran injection, and our measurements were consistent with a mixed endosomal population (pH ∼5.75). Animals injected with dextran sensors s.c. and imaged 24h later had no detectable fluorescence in the vasculature, but meningeal phagocytes showed a marked perinuclear dextran distribution, indicating LE/Ly colocalization, as classically seen in macrophage cell models^15,25^. This was expected, since large dextran polymers (>40 kDa) –composed of glucose units– cannot be digested by lysosomal hydrolases, resulting in accumulation in LE/Lys over time. Our pH measurements demonstrate a marked heterogeneity in LE/Ly acidification (ranging between pH 4.75-6.5). We also found no significant effect of phagocyte proximity to arterioles or venules, or of sex, on LE/Ly acidification. In the dural and subdural spaces we encounter a very heterogeneous macrophage population, expressing variable levels of MHC II and thus differentially involved in antigen presentation^41^. Antigen presenting cells such as dendritic cells and B lymphocytes often possess alkaline lysosomes, which maximizes antigen presentation^42,43^. Imaged dendritic cells and macrophages expressing high MHC II levels may contain incompletely acidified LE/Ly compartments, whereas cells with lower MHC II expression may acidify LE/Lys more efficiently, thus consistent with the heterogeneity in LE/Ly acidification observed in the meninges.

An important aspect of this study is the use of ApHID-dextrans to monitor LE/Ly pH dynamically after cerebral ischemia, or during *in vivo* pharmacological manipulation. We monitored changes in LE/Ly pH after ischemic injury in nearby meningeal phagocytes over time by repeated intravital imaging of the same regions. The inflammatory cell composition of the ischemic lesion is very heterogeneous and includes, in addition to local BAMs and microglia, also monocyte-derived macrophages, neutrophils, and other leukocytes infiltrating the ischemic brain^44^. However, our imaging was focused on the meninges surrounding lesioned arterioles. In this compartment, local resident macrophages and dendritic cells are the first responders following ischemic insult^30,44^, whereas lymphocytes and natural killer cells infiltrate much later^44^. We did not observe any morphologically distinct cells –round in morphology, suggesting infiltration– with endocytosed dextrans in significant numbers until days 7 and 11 post-lesion. This suggests that a significant fraction of cells undergoing LE/Ly acidification might correspond to resident macrophages and/or phagocytes recruited from nearby regions after ischemia, and likely remaining present throughout imaging sessions. Using ApHID we observed a progressive acidification of LE/Ly compartments in these phagocytes, which reached a plateau at 7-11 days, coinciding with the presence of dextran-loaded cell infiltrates. Moreover, using ApHID we were also able to detect LE/Ly alkalinization in meningeal phagocytes, induced by acute chloroquine treatment. Overall, these results demonstrate that ApHID-dextrans can be used to monitor changes in LE/Ly pH dynamically and in real time in both the intact and lesioned brain. Our findings suggest the possibility that pharmacological acidification of LE/Ly compartments in the measured cell types could be explored as a therapeutic strategy to enhance proteolytic activity.

Amino dextrans are affordable and widely available markers, and can be labeled with any fluorescent probes provided they carry an NHS ester. The polymers can be used to label LE/Lys in any organs and animal models. Although we studied the meninges, other organs in the abdomen as well as skin, mammary glands, lungs (reviewed by Jacquemin et al. ^45^) and lymph nodes^46^ can be visualized intravitally by chronic window implantation and are not restricted by the BBB, making our methodology portable to other biological and disease contexts. For these reasons, we believe our approach can be widely implemented and used not only to quantify LE/Ly pH, but also to study endosomal dynamics in various phagocytic cells in homeostatic conditions and disease contexts.

## LIMITATIONS OF THE STUDY

Cellular acidic compartments can be imaged in vivo using nanosensors^40^ or genetically encoded biosensors ^47,48^. The latter ensures selective lysosomal localization but requires expressing the reporters in cell culture or in animal models. With our approach, a fraction of dextran polymers will accumulate in late endosomes and report pH from a somewhat mixed endolysosomal population (lysosomes are constantly fusing with late endosomes in a process mediated by endocytic Ca^2+^ ^49^). Therefore, for applications requiring purely lysosomal sensing, genetically encoded reporters may be preferable.

In our experiments, fluorescence ratios were translated to pH by interpolation to a ratio-to-pH standard curve prepared in fixed J774A.1 macrophages incubated in pH 5.0 buffer. To generate the calibrations, J774A.1 macrophages were loaded with the same dextrans used to measure pH intravitally, chased for 4h to ensure LE/Ly localization, fixed in PFA, and imaged in pH 5.0 buffer at 37 °C using the same acquisition settings applied during intravital imaging. The full ratio-to-pH titration was then reconstructed, anchoring the curve to the measured pH 5.0 ratio and rebuilding the remaining values using complete titration data obtained previously in fixed J774A.1 macrophages. Several lines of evidence support the validity of this approach.

ApHID fluorescence is not significantly altered by 48h incubation in macrophage lysosomes or by PFA fixation, and is preserved in the presence of reactive oxygen species and across various CaCl_2_ and MgCl_2_ concentrations^15^. Moreover, buffer penetration across fixed membranes is maximized using membrane-permeant equilibrators, and the resulting ratio-to-pH calibrations in fixed cells are virtually identical to those obtained in solution using a plate reader, indicating optimal buffer equilibration^15^. Our technical approaches yielded very stable pH values across all imaged animals and technical replicates. Nevertheless, the optimal calibration setup for in vivo measurements would have involved infusing pH 4.0-6.0 buffers into preserved mouse brains under anesthesia, followed by intravital imaging at 37 °C without fixative – an experiment that is not technically feasible.

Regarding ApHID’s dynamic range, the probe has limited discriminating power when measuring pH >6.0. Translated pH values above 6.0 constituted only a small fraction of all LE/Lys measured, but were obtained by extrapolation to the standard calibration and should be interpreted with caution. Nevertheless, in the ischemic lesion experiment, the ratio-to-pH standard curve included pH 4.5-6.5 fluorescence ratio values, yet the interpolated baseline LE/Ly pH values obtained prior to photothrombosis were identical to those of all other ratiometric pH measurements (with calibrations including pH 4.0-6.0 fluorescence ratio values) in meningeal phagocytes (Fig. 3-5 and 7).

Other limitations should be acknowledged. First, transient photothrombotic lesions of single penetrating arterioles differ from larger ischemic models such as middle cerebral artery occlusion in scale and kinetics, and the infiltration dynamics we infer should not be extrapolated directly. Second, two-photon imaging through a cranial window has finite depth penetration, and cranial window surgery itself can induce a degree of meningeal inflammation that we cannot fully exclude as a baseline contributor. Third, while our meningeal phagocyte cell-identity determinations are well supported by fate-mapping and transcriptomic data from the literature^19,20^, definitive assignment of imaged cells to specific macrophage subsets will require co-labeling strategies in future work.

## Supporting information

Supplementary Section

## ACKNOWLEDGEMENTS

This work was supported by NIH grants RF1-AG078244 and R01-HL093324 and the Cure Alzheimer’s Foundation grant CAF-211540-02. S.S.D. was supported by the Swedish Research Council International Postdoctoral Fellowship number 637-2013-503 / D0050301 and the Leon Levy Foundation Fellowship in Neuroscience. M.I. was supported by a NIH Medical Scientist Training Program grant from the National Institute of General Medical Sciences under award number T32GM152349 to the Weill Cornell/Rockefeller/Sloan Kettering Tri-Institutional MD-PhD Program. The authors are grateful to the Weill Cornell Chemistry Core for synthesizing ApHID.

## AUTHOR CONTRIBUTIONS

Conceptualization, S.S.D., F.R.M., J.A., C.I., A.P.; Methodology, S.S.D., S.J.A., M.I., L.F., C.J., J.P., L.M.J.; Validation, S.S.D., S.J.A., M.I., C.J., J.P.; Investigation, S.S.D., S.J.A., M.I., C.J., J.P., E.C.Z.; Resources, F.R.M., S.S.D., J.D.W., J.A., C.I.; Data Curation, S.S.D., S.J.A., L.M.J.; Writing – Original Draft, S.S.D.; Writing – Review & Editing, S.S.D., F.R.M., L.M.J., A.P., M.I., J.D.W., J.A., C.I., E.C.Z.; Visualization, S.S.D., S.J.A., C.J., J.P.; Supervision, S.S.D., F.R.M., A.P., J.A., C.I.; Project Administration, S.S.D., F.R.M.; Funding Acquisition: F.R.M., S.S.D.

## METHODS SECTION

### Experimental Model and Study Participant Details

#### J774A.1 murine macrophage cell culture

J774A.1 murine macrophages (ATCC TIB-67) were grown in Dulbecco’s Modified Eagle’s Medium (DMEM) containing 4.5 g/L glucose and 1 mM sodium pyruvate (Corning 15-013-CV), with 10% fetal bovine serum (FBS, Gemini BenchMark FBS 100-106), 4 mM L-glutamine (Gibco 25030081), and 1% penicillin-streptomycin (Thermo Scientific 15140163) in an incubator at 37 °C with humidified atmosphere and 5% CO_2_. Cells were passed at a subcultivation ratio of 1:5 every 2-3 days.

#### Mouse models

CX3CR1-eGFP adult mice (Jackson Laboratories, stock# 005582) were used to study meningeal phagocyte dextran uptake in vivo; CX3CR1-Lyve1-tdTomato mice (Jackson Laboratories, stock# 033319) were used to monitor perivascular border-associated macrophage dextran uptake, and wild-type adult mice (Jackson Laboratories, stock# 000664) were used to measure LE/Ly pH in meningeal phagocytes. All mice were maintained on a C57BL/6J background. Mice were housed for harem breeding when necessary (one male, two females) and maintained in 12-h dark/light cycle sterile ventilated cages with access to food and water *ad libitum* at Weill Cornell Medicine animal facilities. All animal experiments were conducted in compliance with the Institutional Animal Care and Use Committee of Weill Cornell Medicine.

## Method Details

### Preparation of reagents and dextrans

- *Dextran derivatization with fluorophores.* Polymers of 70 kDa amino-dextrans (Thermo Fisher D1862) or 500 kDa amino-dextrans (Thermo Fisher D7144) were solubilized at 25 or 20 mg/mL in sterile 0.1 M NaHCO_3_ buffer adjusted to pH 8.3. 70 kDa amino dextrans were reacted at polymer:dye 1:4 molar ratio with the following *N*-hydroxysuccinimidyl esters (NHS): NHS-ApHID (custom-made), NHS-5/6-carboxyfluorescein (NHS-fluorescein, Thermo Fisher 46410) and NHS-Alexa 405 (Thermo Fisher A30000). 500 kDa amino dextrans were used to label brain vasculature and were reacted at polymer:dye 1:10 molar ratio with NHS-fluorescein and NHS-Alexa 546, and at 1:22 molar ratio with NHS-Alexa 405. The polymers were incubated with the various NHS esters for 2h at room temperature (RT) with constant rotation. Following reaction, dextrans were purified by extensive dialysis in 20 kDa cutoff Side-A-Lyzer dialysis cassettes (Thermo Fisher 66003) against 1X PBS and stored at 4 °C until use. Dye incorporation was verified by measuring absorbance of the labeled dextrans using a spectrophotometer. When measuring ApHID incorporation, dextrans were diluted in 25 mM citric acid, 25 mM sodium citrate buffer adjusted to pH 3.0 and absorbance was read at 502 nm. When measuring labeling with fluorescein, Alexa 546 and Alexa 405, the dextrans were diluted in pH 7.4 PBS buffer, and absorbance was read at 490, 573 nm and 405 nm.
- *Buffers for measurements in fixed J774A.1 macrophages.* Fixed cells (described below) were incubated in buffers with pH adjusted between 4.0 and 6.5. To prepare the buffers, buffer salts were added to 0.13X PBS as follows: pH 4.0 to 5.5: 50 mM TRIS-maleate; pH 6.0-6.5 buffer: 50 mM sodium phosphate monobasic anhydrous. The buffers were supplemented with 40 mM sodium acetate (Sigma-Aldrich, S8750) and 40 mM methylamine hydrochloride (Sigma-Aldrich, M0505) as membrane-permeant equilibrators (to ensure buffer equilibration across membranes).The pH was determined using an Orion Star A211 pH meter (Thermo Fisher), and acidity was adjusted by adding 1-3 N HCl or 1-10 N NaOH dropwise. The resulting buffer ionic strength of the solutions was approximately 150 mM. After preparation, the solutions were filtered through a 0.4 µm filter membrane and stored at 4 °C. Immediately prior to cell incubation (see below), 40 µM monensin sodium salt (Sigma-Aldrich, M5273) was added to the buffers and aliquots were warmed up to 37 °C.

### Labeling of J774A.1 macrophage late endosomes and lysosomes with 70 kDa dextrans

J774A.1 macrophages were seeded in 35-mm dishes with central 7 mm diameter glass bottom imaging chambers coated with poly-D-lysine at 40,000 cells per chamber and allowed to settle for 1-2h in an incubator at 37 °C under 5% CO_2_. Once settled, cells were incubated with 70 kDa ApHID-, fluorescein-, ApHID-Alexa 546 or ApHID-Alexa 405 amino-dextrans at 0.5 mg/mL in complete DMEM medium overnight at 37 °C, followed by a 3-4h chase in fresh media the following morning to ensure late endosomal and lysosomal localization. Cells were thereafter washed 2X in sterile 1X PBS and fixed in 0.5% PFA (Electron Microscopy Sciences, 15714-S) at 22-25 °C for 5 min followed by 3X wash in 1X PBS. Fixed cells were stored at 22-25 °C protected from light and imaged by two-photon microscopy on the same day.

### Qualitative two-photon excitation spectrum measurements for ApHID-dextrans

Fixed J774A.1 macrophages seeded in 35-mm dishes previously labeled with 70 kDa ApHID-dextrans were incubated in 50 mM TRIS maleate pH 5.5 buffer containing 40 mM sodium acetate, 40 mM methylamine and 40 µM monensin for 20 min at 37 °C. Dishes were then placed on a warm metal plate maintained at 37 °C, allowed to re equilibrate for 5 min and imaged with a two-photon SP8 scope (Leica). The experiment was repeated twice and a total of 2 dishes were imaged.

### Photostability studies in fixed J774A.1 macrophages using two-photon excitation

Fixed J774A.1 macrophages seeded in 35-mm dishes previously labeled with 70 kDa ApHID- or fluorescein-dextrans were incubated in TRIS maleate pH 5.5 buffer for 30 min at 37 °C inside a cell incubator. The buffers were supplemented with 40 mM methylamine hydrochloride, 40 mM sodium acetate and 40 µM monensin for 30 min at 37 °C. Following buffer equilibration, cells were placed on a warm metal plate maintained at 37 °C, allowed to re equilibrate for 5 min and imaged by two-photon microscopy. The experiment was repeated twice. 3-4 dishes per fluorophore condition were imaged, and 3 fields were acquired for each dish.

### Fluorescence ratios-to-buffer pH calibration in fixed J774A.1 macrophages

Fixed J774A.1 macrophages seeded in 35-mm dishes previously labeled with 70 kDa ApHID-Alexa 546 or ApHID-Alexa 405 amino-dextrans were incubated in 2 mL of buffers with pH adjusted to 4.0, 4.5, 5.0, 5.5, 6.0 or 6.5 (see main Methods section), supplemented with 40 mM methylamine hydrochloride, 40 mM sodium acetate and 40 µM monensin at 37 °C prior to addition. The cells were allowed to equilibrate in the buffers for 15-20 min (pH 4.0 and pH 4.5) or at least 30 min (pH 5.0-6.5). Following incubation, the glass coverslips adhered to the 35-mm dishes were quickly detached using a razorblade and carefully mounted over 10 µL of corresponding buffer on a microscopy glass slide (Fisher Scientific, 22-037-246) previously warmed up to 37 °C. The slide was then placed on a warm metal plate maintained at 37 °C and allowed to re equilibrate for 5 min, followed by two-photon microscopy imaging. One coverslip was imaged for each buffer condition, and 4-8 fields were acquired per coverslip. The titrations were repeated 2-3 times.

### Cranial window implantation

Chronic optical access was established with a glass-covered cranial window, as described previously^50^. Mice received dexamethasone (0.2 mg/Kg, s.c.) at least 2h prior to surgery to reduce brain swelling. Bupivacaine (50 µl of 0.5% solution) was given locally at the incision site. Anesthesia was maintained with isoflurane (1.5– 2% in room air) while body temperature was held at 37 °C using a feedback-controlled heating blanket (SomnoSuite with RightTemp, Kent Scientific). Eye ointment was applied in order to prevent eye drying during the procedure. After removal of the scalp and periosteum, a custom titanium headpost was secured over the somatosensory cortex using dental cement (C&B Metabond, Parkell). A 6-mm bilateral craniotomy was made over the parietal cortex, and the exposed brain was sealed with an 8-mm circular borosilicate glass coverslip (Electron Microscopy Sciences, 72296-08), with sterile saline filling the space beneath, and affixed using cyanoacrylate and tissue adhesive (3M, 1469SB). At the end of surgery, mice received buprenorphine (0.5 mg/Kg, s.c.) for postoperative analgesia. Postoperative care included dexamethasone and buprenorphine for three days, and animals were allowed to recover for at least 3 weeks before imaging experiments.

### Systemic delivery of dextran sensors

Fluorescently labeled dextrans were delivered systemically by intraperitoneal (i.p.) or subcutaneous (s.c.) injection. Dextran conjugates were prepared in sterile saline or PBS and passed through a 0.22 µm filter prior to injection. Mice were briefly anesthetized with 3% isoflurane immediately before injection, rather than manually restrained, to minimize the risk of dislodging the chronic cranial window implant. For i.p. delivery, 200–300 µL of dextran solution was injected into the lower left abdominal quadrant using a 27–30 G needle, avoiding the midline and bladder. For s.c. delivery, 100–200 µL was injected into the loose skin over the dorsal interscapular region. Animals recovered from anesthesia within minutes and were returned to their home cages and monitored until imaging. For experiments requiring labeling of late endosomal and lysosomal compartments, dextrans were injected 24 h prior to imaging. The injection site was inspected before each imaging session to confirm complete absorption, defined as absence of residual subcutaneous fluid or bulge at the injection site. For endosomal pH measurements, dextrans were injected 5-10 min prior to imaging and polymer incorporation into vasculature and acidic vesicles was monitored in real time.

### Two-photon microscopy

- *Two-photon excitation spectra recordings for ApHID-dextran.* Imaging was performed using SP8 two-photon microscope (Leica) equipped with a 25× water-immersion objective (1.0 NA) with motorized correction collar. Microscope control and image acquisition were managed with the Leica Application Suite X (LASX, v1.4.7.28982; Leica). Dishes with fixed J774A.1 macrophages were placed on a warm metal plate maintained at 37 °C during image acquisition. ApHID was excited using a two-photon Mai Tai® HP DeepSee tunable laser (MKS – Spectra-Physics). Images were acquired between 690 to 1040 nm excitations with 10 nm wavelength increments. Stacks of images (512 × 512 pixels per frame, 1 µm step size) were acquired for each field. Fluorescence was detected using a high-sensitivity Leica HyD detector with a spectral window adjusted to 500-550 nm. Resonant acquisition mode at 8K Hz was used. Channel gain settings were optimized during initial stack acquisition and kept constant for all subsequent imaging.
- *Photobleaching study in fixed cells.* Imaging was performed using an SP8 two-photon microscope (Leica) equipped with a 25× water-immersion objective (1.0 NA) with motorized correction collar. Microscope control and image acquisition were managed with the Leica Application Suite X (LASX, v1.4.7.28982; Leica). Dishes with fixed J774A.1 macrophages were placed on a warm metal plate maintained at 37 °C during image acquisition. ApHID and fluorescein-dextrans were excited using a two-photon Mai Tai® HP DeepSee tunable laser (MKS – Spectra-Physics) adjusted to 940 nm with the output adjusted to yield 50 mW power at the front element of the objective using an external power meter, as determined with an external laser power meter. Stacks of images (512 × 512 pixels per frame, 1 µm step size, 50 frames per stack, 0.26 s acquisition per frame) were acquired for a total of 30 irradiation cycles. Pixel dwell time was 72 ns. Fluorescence was detected using a high-sensitivity Leica HyD detector with a spectral window adjusted to collect light at 500-550 nm. Resonant acquisition mode at 8K Hz was used. Channel gain settings were optimized during initial stack acquisition and kept constant for all subsequent imaging.
- *Ratiometric imaging of fixed J774A.1 cells in pH-adjusted buffers.* Imaging was performed using an SP8 two-photon microscope (Leica) equipped with a 25× water-immersion objective (1.0 NA) with motorized correction collar. Microscope control and image acquisition were managed with the Leica Application Suite X (LASX, v1.4.7.28982; Leica). Following incubation in pH-adjusted buffers, glass coverslips with fixed J774A.1 macrophages were mounted on microscopy glass slides, placed on a warm metal plate maintained at 37 °C and imaged. ApHID, Alexa 546 and Alexa 405 were excited using a two-photon Mai Tai® HP DeepSee tunable laser (MKS – Spectra-Physics) adjusted to 940 (ApHID-Alexa 546 pair) or 810 nm (ApHID-Alexa 405 pair). Stacks of images (512 × 512 pixels per frame, 1 µm step size) were acquired per field. Fluorescence was detected using adjustable high-sensitivity Leica HyD detectors with spectral windows set to collect light for each probe (Alexa 405 at 415-485 nm; ApHID at 500-550 nm and Alexa 546 at 560-610 nm). Channel gain settings were optimized during initial stack acquisition and kept constant for all subsequent imaging in different buffers.
- *Intravital imaging of dextran incorporation into meningeal phagocytes.* Imaging was performed using an SP8 two-photon microscope (Leica) equipped with a 25× water-immersion as described above. To monitor dextran uptake by meningeal phagocytes in wild-type mice, 4 mice (1 male and 3 females) were imaged prior to, or 24h after injection of 70 kDa ApHID-Alexa 546 dextrans (20 mg/mL; 200-300 µL delivered s.c.). Mice were anesthetized with isoflurane (1-1.5%) and placed on a warm metal plate maintained at 37 °C. 4-10 regions were located and acquired. ApHID and Alexa 546 were excited using a two-photon Mai Tai® HP DeepSee tunable laser (MKS – Spectra-Physics) adjusted to 940. Stacks of images (512 × 512 pixels per frame, 1 µm step size) were acquired per region down to 100 microns in depth, encompassing the dural and subdural meningeal compartments. The day before dextran injection, regions populated with meningeal phagocytes were identified visually by their intracellular autofluorescence. Laser power was maintained constant throughout acquisition sessions (before and after dextran injections). Nile red beads were imaged at the end of each imaging session (Spherotech, 6 microns) in order to control for small day-over-day variations in laser intensity output and detector performance. Fluorescence intensities were normalized to bead intensity to allow for dextran uptake comparison across imaging sessions and animals. To monitor dextran uptake by tdTomato-tagged dural perivascular macrophages, 4 CX3CR1-Lyve1-tdTomato mice (3 males and 1 female) were injected with 70 kDa ApHID-Alexa 405 dextrans (25 mg/mL, 100-300 µL delivered s.c.) and imaged 24h later. Imaging was performed as described above. 5 to 20 regions encompassing the dural compartment were acquired per animal. Fluorescence was detected using adjustable high-sensitivity Leica HyD detectors with spectral windows set to collect light for each probe (Alexa 405 415-485 nm; ApHID at 500-550 nm and Alexa 546 at 560-610 nm). Channel gain settings were optimized during initial stack acquisition and kept constant for all subsequent imaging sessions.
- *Intravital pH imaging of endosomal compartments in meningeal phagocytes.* Imaging was performed using an SP8 two-photon microscope (Leica) equipped with a 25× water-immersion objective as described above. 70 kDa fluorescein-Alexa 546 dextrans were injected in one wild-type male (12 mg/mL; 240 µL i.p.) whereas ApHID-Alexa 546 dextrans were injected in three wild-type males (20 mg/mL; 240 µL i.p.). Immediately following injections, mice were anesthetized with isoflurane (1-1.5%) and placed on a warm metal plate maintained at 37 °C. 4-6 regions with clearly identifiable meningeal macrophages and vasculature were repeatedly acquired over the course of 2h to monitor dextran incorporation into these elements over time. ApHID, fluorescein and Alexa 546 were excited using a two-photon Mai Tai® HP DeepSee tunable laser (MKS – Spectra-Physics) adjusted to 940. Stacks of images (512 × 512 pixels per frame, 1 µm step size) were acquired per region down to 100 microns in depth, encompassing the dural and subdural compartments. Fluorescence was detected using adjustable high-sensitivity Leica HyD detectors with spectral windows set to collect light for each probe (ApHID at 500-550 nm and Alexa 546 at 560-610 nm Red and green channel gain settings were optimized during initial stack acquisition and kept constant for all subsequent imaging, including ApHID calibration stacks performed on the same day.
- *Intravital pH imaging of LE/Ly compartments in meningeal phagocytes.* Imaging was performed using an SP8 two-photon microscope (Leica) equipped with a 25× water-immersion objective as described above. For LE/Ly pH imaging, 70 kDa ApHID-Alexa 546 dextrans were injected in 4 wild-type males and 4 females (20 mg/mL; 200-300 µL s.c.) and imaged 24h later. Next day, mice were injected with 500 kDa fluorescein-Alexa 546 dextrans (20 mg/mL; 400-500 µL i.p.) in order to label vasculature, anesthetized with isoflurane (1-1.5%) and placed on a warm metal plate maintained at 37 °C. 5-6 regions located in the dural and subdural compartments were acquired as described earlier. To study the effects of acute chloroquine diphosphate treatments, 4 mice (one male and three females) were imaged intravitally (as described above) prior to, and 2 and 8 min following acute chloroquine diphosphate delivery (500 mg/Kg; i.p.). 500 kDa fluorescein-Alexa 546 dextrans were injected (20 mg/mL; 500 µL i.p.) to label vasculature. 4 matched regions were acquired in the dura, for each mouse, before and after CQ injection. Red and green channel gain settings were optimized during initial stack acquisition and kept constant for all subsequent imaging, including ApHID calibration stacks performed on the same day
- *Intravital LE/Ly pH imaging of meningeal phagocytes in proximity to arterioles or venules.* Imaging was performed using a Fluoview two-photon microscope (Olympus) equipped with a 25× water-immersion objective (1.05 NA) mounted on a piezo stage (Physik Instrumente P-725K085,) and an OPO laser (Spectra-Physics, InSight DS+;). Microscope control and image acquisition were managed with Fluoview software (FV31S-SW, v2.3.1.163; Olympus). Wild-type mice were injected with ApHID-Alexa 546 dextrans (20 mg/mL; 200-300 µL s.c.) and allowed to recover for 24h. The next day, mice were anesthetized with isoflurane (1.5-2%) and placed on a feedback-controlled heating blanket (SomnoSuite with RightTemp, Kent Scientific) and injected with 500 kDa Alexa 405-dextran retro-orbitally (r.o.) to label vasculature. ). Initial imaging was performed at 940 nm to locate and characterize vascular segments. For reference mapping, low-magnification (5×) scans were acquired to identify imaging regions and vessels. Cells in proximity to dural arterioles or venules (within 1-3 µm distance) were identified based on the vascular blood flow dynamics of their nearest vessel. Penetrating arterioles were distinguished by the presence of vasomotion^27^ and inward blood flow into cortical parenchyma^28^, whereas venules had larger lumens and drained toward the cortical surface. High-resolution vascular movies and image stacks (512 × 512 pixels over a 339.41 × 339.41 µm field, 1 µm step size) were acquired. 4-5 regions were acquired in the dural compartment for each mouse. Detection of fluorescence was achieved through following band pass filters: Red (Alexa 546; 575-645 nm), Green (ApHID; 495-540 nm), Blue (Alexa 405; 410-460 nm). Red and green channel gain settings were optimized during initial stack acquisition and kept constant for all subsequent imaging, including ApHID calibration stacks performed on the same day.
- *Intravital LE/Ly pH imaging of meningeal phagocytes in regions subjected to photothrombosis-induced ischemic lesion.* Imaging was performed on a Fluoview two-photon microscope (Olympus) equipped with a 25× water-immersion objective as described above. 4 mice (two males and two females) were studied. The animals were injected with 70 kDa ApHID-Alexa 546 dextrans (20 mg/mL; 200 µL s.c.) the day before each imaging session. Animals were anesthetized with isoflurane (1.5–2%) and received a retro-orbital injection of Rose Bengal (20 mg/ml in PBS, 0.1 ml) prior to imaging. Detection of fluorescence was achieved through following band pass filters: Red (Alexa 546; 575-645 nm), Green (ApHID; 495-540 nm), Blue (Alexa 405; 410-460 nm). Initial imaging was performed at 940 nm to locate and characterize vascular segments. For reference mapping, low-magnification (5×) scans were acquired to identify imaging regions and mark the site for subsequent photothrombotic induction. The objective was then switched to 25× and the field of view centered on the target arteriole (12–16 µm diameter). Penetrating arterioles were selected for photothrombosis, and identified by the presence of vasomotion and inward blood flow as described above. High-resolution vascular movies and image stacks (512 × 512 pixels over a 339.41 × 339.41 µm field, 1 µm step size) were acquired. Red and green channel gain settings were optimized during initial stack acquisition and kept constant for all subsequent imaging, including ApHID calibration stacks performed on the same day. For photothrombosis induction, the field of view was zoomed to include only the lumen of the target arteriole. The excitation wavelength was switched to 720 nm, and the vessel was scanned at high frame rates (30–60 Hz, depending on area) while monitoring images in real time^32^. The scanning was terminated when blood clots were formed. The laser wavelength was then returned to 940 nm, and a post-occlusion image stack was acquired at the same site as the baseline stack. To assess vascular and tissue responses over time, follow-up imaging was performed at 1, 4, 7, and 11 days after clot induction. Vasculature was labeled by injection of 500 kDa amino dextrans labeled with Alexa 405 NHS ester (20 mg/mL; 100 µl r.o.) prior to imaging. For each session, the same field of view was relocated using the structural reference map acquired before stroke induction, ensuring consistent registration across time points. Image stacks were then collected to evaluate tissue changes surrounding the targeted arteriole. At day 1, residual Rose Bengal leakage following ischemic lesion interfered with the Alexa 546 signal, preventing accurate estimation of pH immediately after laser illumination. However, fields could be imaged after Rose Bengal injection, prior to laser illumination-clotting, and were used to quantify pH at time 0 (pre-lesion). Following laser illumination-clotting, subsequent imaging confirmed that all targeted vessels had been reperfused.
- *Intravital imaging of fixed J774A.1 macrophages for ratio-to-pH calibration.* Fixed J774A.1 macrophages loaded with the same ApHID-Alexa 546 or ApHID-Alexa 405 dextrans injected to mice were incubated in pH 4.0-6.5 buffers at 37 °C for 10 (pH 4.0 and pH 4.5) to 30 min (pH 5.0-6.5) and imaged as described in “*Ratiometric imaging of fixed J774A.1 cells in pH-adjusted buffers* “. The same blue, green and red channel settings used during intravital imaging were applied. To generate full standard ratio-to-pH calibrations anchored in a single pH 5.0 calibration point, ApHID/Alexa 546 and ApHID/Alexa 405 ratios corresponding to pH 5.0 in fixed cells (imaged at the end of each intravital imaging session) were calculated and used to reconstruct fluorescence ratio-to-pH calibrations based on complete titrations (including pH 4.0 to pH 6.5 ratios) prepared previously in fixed J774A.1 macrophages imaged at 37 °C using the same two-photon microscope.
- *Intravital imaging of meningeal and parenchymal vasculature loaded with dextrans.* Imaging was performed using an SP8 two-photon microscope (Leica) equipped with a 25× water-immersion objective as described above. 2 wild-type female mice were injected with 70 kDa Alexa 405-fluorescein-dextrans (20mg/mL; 400 µL i.p.) whereas 3 mice (1 male and two females) were injected with 500 kDa fluorescein-Alexa 546 dextrans (20 mg/mL; 400 µL i.p.). Mice were anesthetized with isoflurane (1-1.5%) and placed on a warm metal plate maintained at 37 °C. Alexa 405 and fluorescein were excited using a two-photon Mai Tai® HP DeepSee tunable laser (MKS – Spectra-Physics) adjusted to 810 nm, whereas ApHID-Alexa 546 dextrans were excited at 940 nm. In mice injected with fluorescein-Alexa 405 dextrans, two additional channels (far red and green channels excited at 910 nm) were acquired. A total of 5 regions (Alexa 405-fluorescein-dextran study) and 22 regions (fluorescein-Alexa 546 dextran study) were imaged. Fluorescence was detected using adjustable high-sensitivity Leica HyD detectors with spectral windows set to collect light for each probe (Alexa 405 at 418-485 nm, ApHID at 500-550 nm, Alexa 546 at 560-610 nm and far red channel at 660-720 nm). Channel gain settings were optimized during initial stack acquisition and kept constant for all subsequent imaging.

### Quantification and Statistical Analysis

#### Digital Image Analysis

Digital image analysis and quantification was done using FIJI (ImageJ) ^51^ version 1.54f for Windows (https://fiji.sc/) and MetaMorph version 6.7.1 for Windows (Molecular Devices, San Jose, California USA, www.moleculardevices.com).

- *Qualitative two-photon ApHID-dextran excitation spectrum.* Stacks of images were saved as Tiff files using FIJI Bioformats, and Metamorph was used to quantify fluorescence. The green fluorescence channel (ApHID) was corrected by subtracting the 5th intensity percentile from each frame in the stacks and the sum projection of each corrected stack (sum of each image in the stack) was generated. ApHID integrated intensity from the sum projections was quantified and plotted against excitation wavelength using GraphPad Prism.
- *Photostability studies in J774A.1 macrophages.* Stacks of images for each acquired field were saved as Tiff files using FIJI Bioformats. The green fluorescence channel (ApHID or fluorescein) was corrected for background intensity by subtracting the 5^th^ percentile intensity value for each image in the stack. A sum projection for each stack was generated, and total integrated intensity (F) per field was measured. Fluorescence was normalized to the intensity corresponding to the first irradiation cycle (F_0_) and plotted as F/F_0_ against cycle time using GraphPad Prism.
- *Ratiometric pH imaging of dextran sensors in fixed J774A.1 macrophages and meningeal phagocytes.* Leica LASX native image stack files were exported to Tiff files using FIJI Bioformats plugin and saved as a separate stack. When quantifying LE/Ly pH in meningeal phagocytes, image stacks were subdivided into smaller sub-stacks containing single or near single cell slices using FIJI. Sub-stacks were then analyzed using Metamorph. Each plane in a sub-stack was corrected for background fluorescence by removing the 10^th^ intensity percentile from each recorded channel. Next, the total sum projection was generated for each corrected sub-stack, and regions of interest (ROI)(s) were drawn around each cell according to their perinuclear dextran distribution, which provided a morphological reference. A threshold was selected and applied to the pH-independent channel (Alexa 546) selecting all intracellular vesicular dextran signal and a binary mask was generated and applied to the pH-sensitive (ApHID) and pH-independent channels. Integrated intensity was then measured for each ROI (corresponding to each dissected meningeal phagocyte) in each masked sub-stack and values were logged on an Excel spreadsheet. ApHID/Alexa 546 intensity ratios were calculated per cell and interpolated to pH using a ratio-to-pH calibration prepared in fixed J774A.1 macrophages, previously labeled with the same dextrans used to label meningeal phagocytes in vivo, but incubated in pH 5.0 buffer in parallel. Stacks of images acquired from fixed macrophages were quantified as described above, but a large ROI was created including all J774A.1 macrophages in the field (as opposed to dissecting individual cells). ApHID/Alexa 546 ratios per field were calculated instead, corresponding to fluorescence ratios at pH 5.0. These ratios were used to reconstruct a ratio-to-pH calibration using complete titration data acquired using a full array of pH-adjusted buffers (pH 4.0, 4.5, 5.0, 5.5 and 6.0) previously acquired with the same two-photon microscope. These titrations were fit to four-parameter sigmoids using GraphPad Prism, with an IC50 (pKa) of ∼5 (corresponding to ApHID’s pKa). ApHID/Alexa 546 ratios calculated for each meningeal phagocyte were translated into pH values by interpolation to the ratio-to-pH calibration using GraphPad Prism.
- *Color-coded ratio image generation based on ApHID-Alexa 546 ratiometric pH imaging.* To generate color-coded, pixel-by-pixel ratiometric pH images, background-corrected stacks (see previous sections above) were thresholded using an intensity threshold applied to the Alexa 546 channel. A mask was then generated, which was applied to both the Alexa 546 and ApHID channels, so that only fluorescence corresponding to acidic compartments was conserved. Next, a Gaussian filter (7x7 pixels) was applied to all images. A 24-bit ratio image was generated with the filtered ApHID and Alexa 546 channels using the “ratiometric image” module in MetaMorph. ApHID/Alexa 546 ratios corresponding to 4.0 or pH 6.0 were assigned as minimum and maximum ratios, respectively. In the ratio images, blue hue indicates mildly acidic pH whereas tones toward green and red correspond to acidic pH.
- *Study of the effect of depth-associated light scattering on fluorescein and Alexa 546 dextrans in labeled vasculature.* Stacks of images composed of 3 channels (blue-green-red channels excited at 940 nm) were analyzed with Metamorph. Stacks were split and corrected for background fluorescence by subtracting the 10th percentile of total signal for each frame in the stack. An intensity threshold was applied to the red channel to dissect lipofuscin granules, which were further discriminated by size using integrated morphology analysis (IMA). Each segmented vesicle was enlarged using the morphological dilate module, and a negative binary mask was generated and applied to the green and red channels to eliminate lipofuscin signal. Remaining fluorescence in these channels was further thresholded to select labeled vasculature and a positive mask was generated and applied to the channels to filter out non-vascular signal. Vasculature in cleaned images was then segmented using the angiogenesis tube formation module applied to the masked fluorescein channel. Finally, integrated intensity was quantified in the cleaned green and red channels. Green/Red fluorescence ratios were calculated for each plane in the stack and averaged for each region imaged and mouse (n=3 mice). Ratios were plotted against depth (z distance 1 micron between individual frames) using GraphPad Prism.

#### Statistical Methods

Statistical analyses were performed using GraphPad Prism version 10.3.1 for Windows (GraphPad Software, Boston, Massachusetts, USA, www.graphpad.com). Extended Data 2 includes all tables with data originating from all measurements carried out for all experiments. Unless stated otherwise, we did not consider mouse sex as a biological variable.

Experiments in solution were repeated independently at least twice, and mean ± SEM are shown (Fig. 1 and Suppl. Fig. 1). For dextran uptake experiments in vivo, 4 CX3CR1-eGFP mice were injected with Alexa 405-dextrans whereas 3 animals were left untreated as control (Fig. 2I). The fraction of cells labeled with dextrans vs. the fraction of unlabeled cells across conditions was compared using a linear mixed effects model with a fixed effect of condition and a random effect of Animal ID due to the repeated measurements taken on each animal across the conditions, followed by Tukey’s multiple comparison test. For dextran uptake by dural perivascular CX3CR1-Lyve1-tdT^+^ macrophage experiments, 4 CX3CR1-Lyve1-tdT mice were injected and imaged (Fig. 2L). The fraction of dextran-labeled vs unlabeled macrophages was compared using a paired Student’s t-test. In plots 2I and 2L, circles and bars represent cell fraction ±SEM.

For ratiometric imaging of endosomal compartments using fluorescein-Alexa 546 dextrans, 1 wild-type mouse was injected with dextrans and imaged. In plot 3E, bars represent mean fluorescein/Alexa 546 fluorescence ratio for meningeal phagocytes, blood vessels or fixed J774A.1 macrophages ±SD whereas geometric shapes correspond to mean fluorescence ratios for individual cells, blood vessels or fixed cell fields, respectively. For ratiometric pH imaging of endosomal compartments using ApHID, 3 mice were injected with ApHID-Alexa 546 dextrans and imaged. In plot 3H, bars indicate mean ApHID/Alexa 546 fluorescence ratio for meningeal phagocytes, blood vessels or fixed J774A.1 macrophages ±SD, whereas geometric shapes indicate values for individual cells, blood vessels or fixed cell fields. In plot 3I, the ApHID/Alexa 546 vs. buffer pH titration shown results from the average of 3 calibrations (see Extended Data 2); circles and bars (fitted within the circles) indicate mean ApHID/Alexa 546 ratio ±SEM whereas the dotted line indicates the accuracy of the fit. In plot 3J, bars indicate mean endosomal pH ±SD for each mouse separately (n=3 mice analyzed) whereas geometric shapes represent endosomal pH per meningeal phagocyte. Plot 3K shows all quantified cells pooled together (dotted lines indicate median, 25^th^ and 75^th^ quartiles).

For quantification of endosomal dextran accumulation in meningeal phagocytes, 4 wild-type mice were injected with dextrans and imaged. In panel 4C3, bars indicate mean integrated intensity ±SEM. Large geometric shapes correspond to mean integrated intensity for individual mice whereas small brown dots indicate values for individually segmented cells, pooled for all mice. For ratiometric pH imaging of LE/Ly compartments in meningeal phagocytes, 8 wild-type mice (4 males and 4 females) were quantified and sex was considered as biological variable. In plot 4G, violin plots integrate LE/Ly pH for all individual meningeal phagocytes quantified (a total of 4093 cells were segmented) sitting either away from vasculature (“NP”), or close proximity to vasculature (“Periv”), or pooled together for all mice. Dotted lines indicate mean, 25^th^ and 75^th^ quartiles. In plot 4H, bars indicate mean LE/Ly per mouse ±SEM whereas geometric shapes correspond to LE/Ly pH for individual mice. Mean LE/Ly was compared between male and female mice, and between cells in close proximity or away from vasculature using a linear mixed effects model with a fixed effect of condition and a random effect of Animal ID due to the repeated measurements taken on each animal across the conditions, followed by Tukey’s multiple comparison test (n.s.; p≥0.05, all comparisons). In plots 4I, geometric shapes and bars indicate mean fraction of cells within a given pH range per mouse ±SEM. Differences in cell fraction mean ranks were assessed using the non-parametric paired Friedman test with multiple comparisons by Dunn’s test.

To measure the effects of meningeal phagocyte proximity to venules or arterioles on LE/Ly pH, 3 male mice were quantified. In plot 5F, bars indicate mean LE/Ly pH associated to proximity to arterioles (A) or venules (V) ±SEM. Mean LE/Ly pH values were compared using the paired Student’s t test.

To measure meningeal phagocyte LE/Ly pH in areas subjected to ischemic lesion, 4 mice (2 males and 2 females) were quantified. In plot 6L, bars indicate mean LE/Ly pH per optical plane and post-lesion time (n=4 mice) ±SEM. In plot 6M, geometric shapes and bars indicate mean LE/Ly pH per lesioned area and post-lesion time (1-2 lesioned areas per mouse, 4 mice, indicated by a specific geometric shape) ± SEM. Mean LE/Ly before or 4, 7 and 11 days post-ischemic lesion was compared using a linear mixed effects model followed by Tukey’s multiple comparison test.

To measure meningeal phagocyte LE/Ly pH following acute CQ treatment, 4 mice were quantified. In plot 7D, bars indicate mean LE/Ly per mouse and post-injection time ±SEM. Large geometric shapes correspond to LE/Ly pH values for individual mice whereas smaller brown circles indicate values for individual fields acquired within mice. Mean LE/Ly pH between post-imaging timepoints were compared using a linear mixed effects model, followed by Tukey’s multiple comparison test.

D’Agostino-Pearson and/or Kolmogorov-Smirnov tests were used to test whether the various data arrays met the assumption of normality. If so, parametric tests were used to compare means between conditions.

Otherwise, the data was transformed or non-parametric tests were used. Forsythe’s and Bartlett’s tests were used to assess differences in the data’s variance between conditions. P-values are shown as p≥0.05 (ns), p<0.05 (*), p<0.01 (**), p<0.001 (***), and p<0.0001 (****)

