## Supplementary Section for "Intravital pH Imaging using the sensor ApHID Reveals Incomplete Lysosomal Acidification in Resident Meningeal Phagocytes"

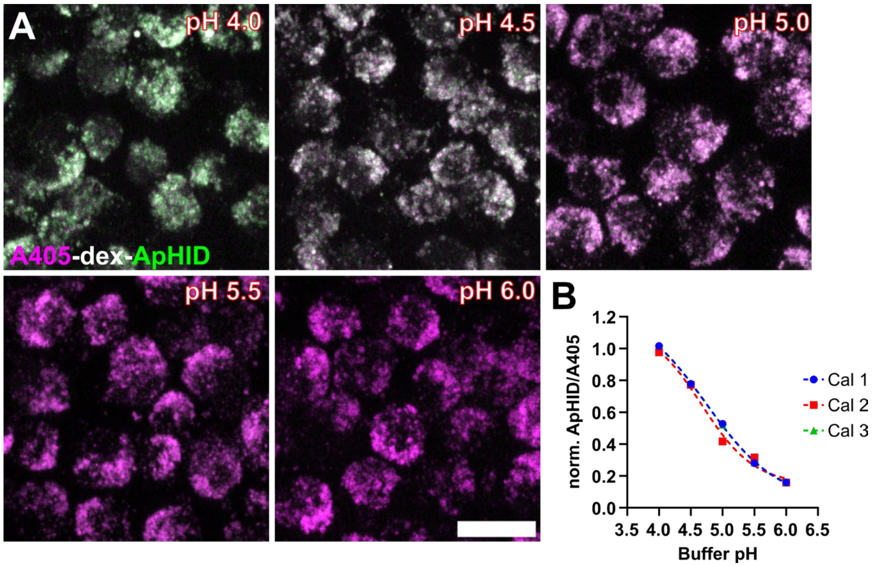


**Supplementary Figure 1.** Dextrans labeled with ApHID and Alexa 405 show pH dependence under two-photon excitation. (A) Fixed J774.1 macrophages labeled with dextrans conjugated with ApHID and Alexa Fluor 405 (pH-independent). Cells were incubated in buffers ranging from pH 4.0 to 6.0, containing membrane-permeant equilibrators, for 10–30 min and imaged at 37°C under two-photon 810 nm excitation. ApHID/Alexa Fluor 405 fluorescence ratios plotted against buffer pH were fit to a four-parameter sigmoidal function, with an $IC_{50}$ corresponding to ApHID's pKa of ~5 (B). The experiment was repeated three times. One dish was imaged per pH condition, and four fields were acquired per dish. Symbols indicate mean fluorescence ratio. Bars (SEM) fit within the symbols. Scale bar: 20 μm.

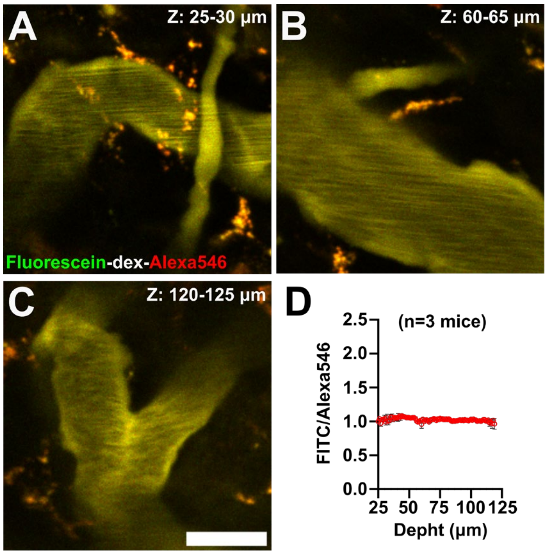


**Supplementary Figure 2. Fluorescein/Alexa 546 dextran fluorescence ratios remain stable within ~100 μm in brain depth. (A-D)** Meningeal vasculature in wild-type mouse brain loaded with 500 kDa fluorescein-Alexa 546 dextrans imaged at various depths (A-C). Fluorescein/Alexa 546 ratios were plotted against brain depth, rendering a lineal fit (D, n=3 mice).

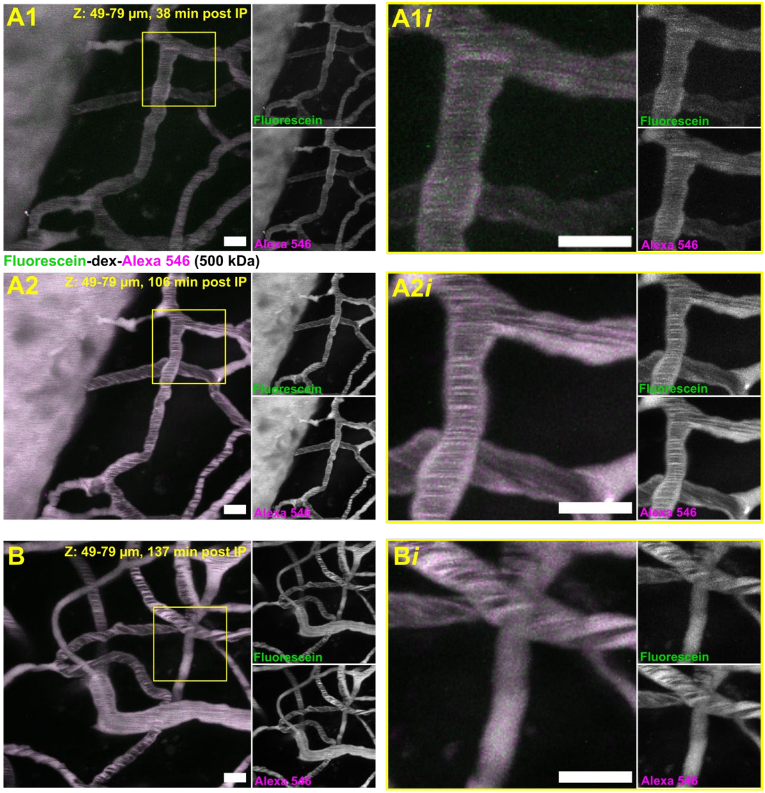


**Supplementary Figure 3. Meningeal phagocytes do not internalize 500 kDa dextrans injected intraperitoneally in significant amounts during intravital imaging. (A)** Intravital imaging of wild-type mouse meninges through a cranial window. Vasculature was labeled by intraperitoneal (IP) injection of 500 kDa dextrans conjugated to fluorescein and Alexa Fluor 546 and imaged 38 min (A1) and 107 min (A2) after injection. The region highlighted by a yellow square in (A1) is shown at higher magnification in (A1i). **(B)** A second field of view with labeled vasculature imaged 137 min after IP dextran injection. The region highlighted by a yellow square in (B) is shown at higher magnification in (Bi). Scale bars: 10 µm.

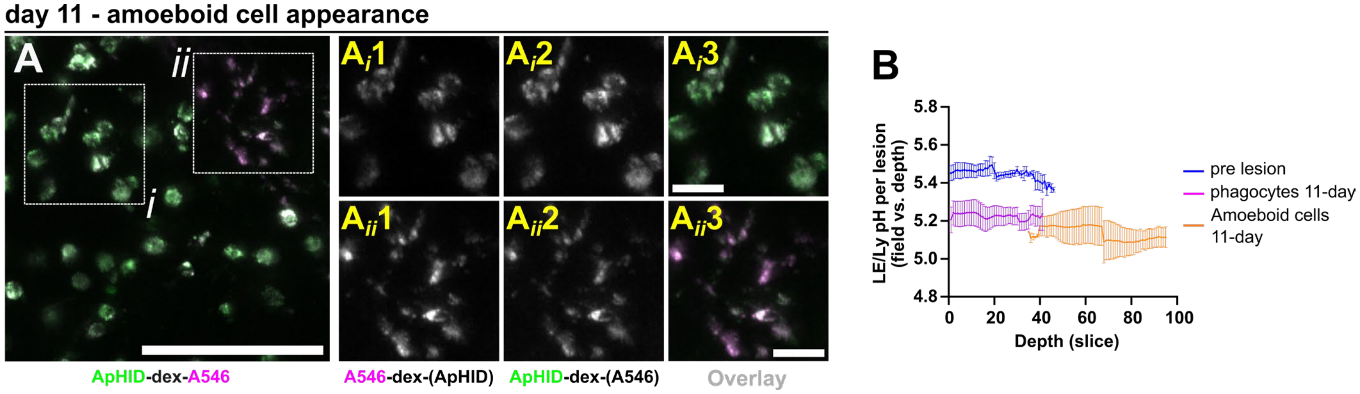


**Supplementary Figure 4. Amoeboid cells infiltrate meningeal regions subjected to photothrombosis 11 days post lesion. (A)** 11 days after photothrombosis, amoeboid phagocytes were observed in deeper cortical layers. Two regions of interest (i, ii; white squares) are shown, with (Ai1–3) highlighting amoeboid cells and (Aii1–3) phagocytes with regular morphologies (A546, ApHID, and overlay). **(B)** Mean meningeal phagocyte LE/Ly pH (1-2 lesioned areas per mouse, 4 mice in total) plotted against depth, including pH of cells with amoeboid morphology (orange lines). Bars indicate mean LE/Ly pH ± SEM. Scale bars: large panels: 100 µm (A); insets: 20 µm (Ai-ii). Abbreviations: A546: Alexa Fluor 546; A405: Alexa Fluor 405.
